# Genome-wide CRISPRi screening identifies OCIAD1 as a prohibitin client and regulatory determinant of mitochondrial Complex III assembly in human cells

**DOI:** 10.1101/2021.02.16.431450

**Authors:** Maxence Le Vasseur, Jonathan R. Friedman, Marco Jost, Jiawei Xu, Justin Yamada, Martin Kampmann, Max A. Horlbeck, Michelle R. Salemi, Brett S. Phinney, Jonathan S. Weissman, Jodi Nunnari

**Affiliations:** Department of Molecular and Cellular Biology, College of Biological Sciences, University of California, Davis, CA, 95616, USA; Department of Cell Biology, University of Texas Southwestern Medical Center, Dallas, TX, 75390, USA; Department of Cellular and Molecular Pharmacology, University of California at San Francisco, San Francisco, CA, 94158, USA; Howard Hughes Medical Institute, University of California at San Francisco, San Francisco, CA, 94158, USA; Department of Microbiology and Immunology, University of California at San Francisco, San Francisco, CA, 94158, USA; Institute for Neurodegenerative Diseases and Department of Biochemistry and Biophysics, University of California at San Francisco, San Francisco, CA, 94158, USA; Chan-Zuckerberg Biohub, San Francisco, CA, 94158, USA; Proteomics Core Facility, University of California, Davis, CA 95616, USA; Whitehead Institute, Cambridge, MA, USA; Department of Biology, Massachusetts Institute of Technology, Cambridge, MA, USA

## Abstract

Dysfunction of the mitochondrial electron transport chain (mETC) is a major cause of human mitochondrial diseases. To identify determinants of mETC function, we screened a genome-wide human CRISPRi library under oxidative metabolic conditions with selective inhibition of mitochondrial Complex III and identified OCIA domain-containing protein 1 (OCIAD1) as a Complex III assembly factor. We find that OCIAD1 is an inner mitochondrial membrane protein that forms a complex with supramolecular prohibitin assemblies. Our data indicate that OCIAD1 is required for maintenance of normal steady state levels of Complex III and the proteolytic processing of the catalytic subunit cytochrome *c_1_* (CYC1). In OCIAD1 depleted mitochondria, unprocessed CYC1 is hemylated and incorporated into Complex III. We propose that OCIAD1 acts as an adaptor within prohibitin assemblies to stabilize and/or chaperone CYC1 and to facilitate its proteolytic processing by the IMMP2L protease.

## Introduction

Mitochondria are double membrane-bound organelles of endosymbiotic origin that produce most of the ATP in eukaryotic cells through oxidative phosphorylation (OXPHOS) (Mitchell, 2011). OXPHOS depends on the mitochondrial electron transport chain (mETC), which transfers electrons from NADH and succinate to molecular oxygen. The mETC is comprised of a series of four large inner mitochondrial membrane (IMM) complexes (CI-CIV) that assemble into supercomplexes of defined stoichiometry (Letts and Sazanov, 2017). Substrate oxidation-driven electron transfer is coupled to the translocation of protons across the IMM to generate an electrochemical gradient harvested by the ATP synthase (CV) for ATP production. In addition, the mETC and the associated tricarboxylic acid (TCA) cycle support a network of metabolic functions. The mETC helps maintain the redox balance of carrier pairs involved in hundreds of biochemical reactions (Luna-Sánchez et al., 2017; Titov et al., 2016; Wang and Hekimi, 2016; Ying, 2008; Ziosi et al., 2017), a basic requisite for sustaining metabolism in living cells, and is also essential for generating the proton gradient that drives the import of nuclear-encoded mitochondrial proteins across the IMM (Eilers et al., 1987; Martin et al., 1991; Pfanner and Neupert, 1986; Schleyer et al., 1982). Perturbing the assembly or function of the mETC can lead to multisystem mitochondrial disorders (Chinnery, 1993; Rodenburg, 2016; Tucker et al., 2013; Wanschers et al., 2014) and is linked to more general pathologies, such as diabetes (Antoun et al., 2015; Ramírez-Camacho et al., 2020), neurodegeneration (Devi et al., 2008; Giachin et al., 2016; Keeney et al., 2006), heart diseases (Andreu et al., 2000; Casademont and Miró, 2002; Hagen et al., 2013; Valnot et al., 1999), and cancer (Hoekstra and Bayley, 2013; Janeway et al., 2011; Pantaleo et al., 2014; Urra et al., 2017; Vranken et al., 2015).

The biogenesis of the mETC requires the concerted expression of nuclear and mitochondrial DNA (mtDNA) encoded genes and is highly regulated. Coordination of mETC subunits of dual origin occurs in part via the formation of modular intermediates within mitochondria that assemble sequentially into functional complexes (Aich et al., 2018; Guerrero-Castillo et al., 2017; Lobo-Jarne et al., 2020; Ndi et al., 2018; Stephan and Ott, 2020; Vranken et al., 2015). Assembly of mETCs strategically occurs in specialized domains that link protein import, membrane insertion, and assembly machineries (Singh et al., 2020; Stoldt et al., 2018). Prohibitins are thought to promote mETC assembly and quality control by assembling into inner membrane ring-like scaffold structures that specify local protein and lipid composition (Nijtmans et al., 2000; Singh et al., 2020). In mammalian cells, prohibitins associate with a variety of inner membrane proteins, including mitochondrial translocases, subunits of mETC, the DnaJ-like chaperone DNACJ19 and m-AAA proteases (Nijtmans et al., 2000; Richter-Dennerlein et al., 2014; Steglich et al., 1999; Yoshinaka et al., 2019). The interaction of prohibitin with these key assembly and quality control proteins either directly modulates their activities and/or influences their client interactions to influence and potentially coordinate a plethora of mitochondrial functions.

Here we use an unbiased genome-wide CRISPRi approach to screen for human genes modulating the cellular response to antimycin A, a chemical inhibitor of mitochondrial Complex III. Complex III, also called ubiquinol-cytochrome *c* oxidoreductase or cytochrome *bc1*, is centrally situated within the mETC. Complex III is an obligate homodimeric enzyme (CIII_2_) embedded in the inner membrane with each monomer composed of 10-11 subunits. Only three subunits contain catalytically active redox groups: cytochrome b (MT-CYB), cytochrome *c_1_* (CYC1), and the Rieske iron-sulfur protein (UQCRFS1), with other accessory subunits that likely stabilize the assembly (Lee et al., 2001; Malaney et al., 1997). We identified OCIAD1, a poorly characterized protein, as a key regulator of Complex III biogenesis. Our data indicate that OCIAD1 is a client of prohibitin supramolecular assemblies and is required for the IMMP2L-dependent proteolytic processing of the catalytic subunit CYC1. Thus, we postulate that within prohibitin assemblies OCIAD1 facilitates CYC1 proteolytic processing by the IMMP2L.

## Results

### Genome-wide CRISPRi screen for antimycin sensitivity identifies Complex III molecular determinants

CRISPR screens have emerged as a powerful approach to identify key genes regulating molecular processes in human cells (Gilbert et al., 2014; Jost et al., 2017; To et al., 2019). To identify regulatory determinants of mitochondrial function, we screened for genes that either sensitized or protected against antimycin A, a selective inhibitor of mitochondrial respiratory Complex III. Candidate genes were identified using a genome-scale CRISPRi screen performed in human K562 cells stably expressing the dCas9-KRAB transcriptional repressor (Gilbert et al., 2013). Cells were infected with the hCRISPRi-v2 sgRNA pooled library containing 10 sgRNAs per gene (Horlbeck et al., 2016) and grown for six days in glucose-free media containing galactose, which favors oxidative metabolism over glycolysis. The cell population was then halved and subjected to four cycles of treatment with either vehicle or antimycin A (3.5-3.75nM; 24h treatment, 48h post-washout recovery), which created a growth difference of approximately 3-4 doublings between treated and untreated cells (Figure 1A). Following the final recovery phase, cells were harvested at ∼750 cells per sgRNA and sgRNA-encoding cassettes were PCR-amplified from genomic DNA. The abundance of each individual sgRNA was then quantified by next-generation sequencing and a phenotype score (ρ) was calculated for each gene as described (Gilbert et al., 2014; Jost et al., 2017; Kampmann et al., 2013). This phenotype score represents the differential pressure each sgRNA exerts on cell growth in the presence versus absence of antimycin A. Positive ρ values indicate protection and negative ρ values indicate sensitization to antimycin A.

**Figure 1.**
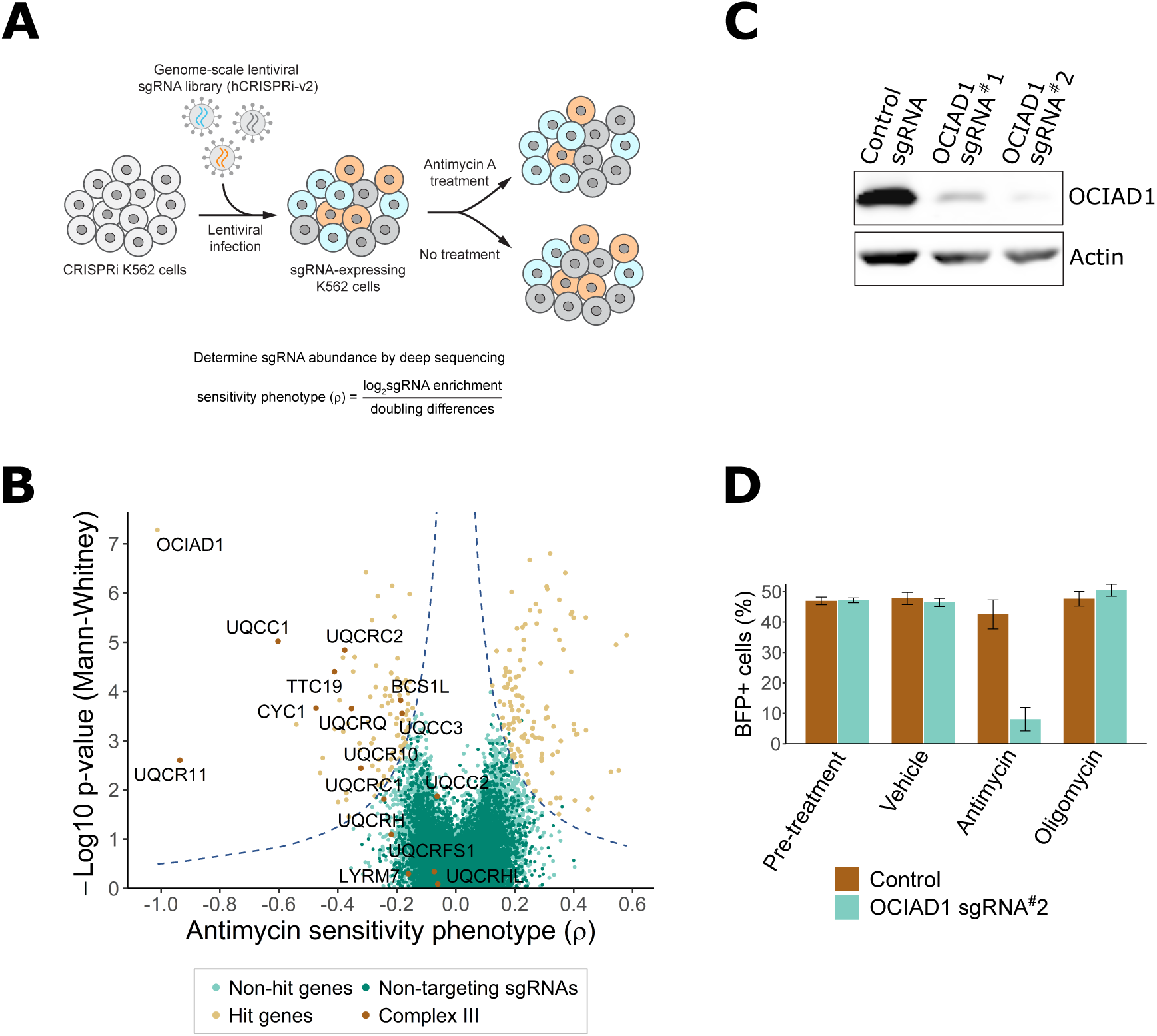
Genome-scale CRISPRi antimycin screen identifies genes regulating mitochondrial Complex III. A) Schematic overview of the genome-wide CRISPRi screen. K562 dCas9 cells stably expressing dCas9-KRAB were infected with a pooled genome-scale sgRNA library. After growth in galactose, cells were subjected to 4 pulses of antimycin A or vehicle treatment followed by a 48h recovery period. After the last antimycin A pulse, genomic DNA from each condition was isolated and sgRNA abundance was quantified by deep sequencing. B) Volcano plot showing the statistical significance (y axis) vs phenotype scores (ρ, x axis) of control non-targeting and genome-wide targeting sgRNAs. Knockdown of Complex III structural proteins and assembly factors sensitized cells to antimycin A. Genes were considered a hit if they scored above a threshold of ρ z-score x -log_10_ p-value of 7 (dashed line). C) CRISPRi knockdown of OCIAD1 expression. Western blot showing the expression level of OCIAD1 in K562 dCas9-KRAB cells stably expressing either a control non-targeting sgRNA or two different sgRNAs against OCIAD1. CRISPRi-based silencing reduced OCIAD1 protein expression by ∼90%. D) Validation of the OCIAD1 phenotype. K562 dCas9 cells were mixed with an equal number of K562 dCas9-KRAB BFP+ cells stably expressing a non-targeting sgRNA (brown bars) or a sgRNA against OCIAD1 (light blue bars). Cell mixtures were then treated with the drug or a vehicle for 24h. The percentage of BFP+ cells in the cell mixtures was measured by flow cytometry before and 24h after treatment. OCIAD1 silencing selectively sensitized cells to antimycin treatment.

Using this approach, we identified 217 genes that significantly modulated sensitivity to antimycin A under oxidative conditions (Figure 1B, source data 1 and 2). Knockdown of 128 of these genes protected against antimycin A. Gene ontology (GO) enrichment analysis performed on this group identified an enrichment of genes encoding for mitochondrial respiratory chain Complex I. Complex I is the most upstream entry point into the electron transport chain and is composed of 44 unique subunits, 37 of which are encoded by the nuclear DNA with the remaining 7 subunits encoded by the mitochondrial genome (Fiedorczuk et al., 2016; Guerrero-Castillo et al., 2017). Knockdown of about one-sixth of the nuclear-encoded Complex I subunits, as well as additional assembly factors, significantly protected against antimycin A treatment (Figure 1-figure supplement 1B,C). Complex I subunit hits were distributed on all Complex I assembly modules except the proximal portion of the peripheral arm, indicating that the protective response is likely dependent on a general loss of Complex I function. Knockdown of genes encoding components of the TCA cycle also protected against antimycin A treatment, including those encoding enzymes that participate in both forward flux through this pathway to maintain oxidative phosphorylation and reverse flux for reductive carboxylation. Other protective hits included a protective gene encoding an assembly factor of Complex II, which connects the TCA cycle to the respiratory chain, upstream of Complex III, as well as genes encoding the mitochondrial pyruvate carrier and pyruvate dehydrogenase, which connects glycolysis with the TCA cycle (Figure 1-figure supplement 1D,E). It was recently reported that the loss of mitochondrial Complex I activity suppressed toxicity caused by oligomycin, an ATP synthase inhibitor, and to a lesser extent by antimycin A, by promoting glycolysis and reductive carboxylation (To et al., 2019). However, the suppressive effect we observe is potentially inconsistent with this mechanism as our screen was performed under different metabolic conditions that promote oxidative metabolism and suppress glycolysis. Thus, it is possible that the mechanism of antimycin A toxicity suppression in our screen was a consequence of a reduction in respiratory chain activity upstream of Complex III to protect against production of ROS, further suggesting that multiple suppressive mechanisms for antimycin toxicity may exist, dependent on cellular metabolic status.

In our screen, knockdown of 89 genes sensitized cells to antimycin A treatment (Figure 1B), including 9 of the 15 nuclear-encoded Complex III subunits or assembly factors (Figure 1-figure supplement 2). Consistent with this, gene ontology enrichment analysis identified Complex III as the most enriched term for antimycin A toxicity (Figure 1-figure supplement 2A). These data validate the screen and confirm that the mechanism of growth inhibition by antimycin A was a consequence of Complex III inhibition.

In addition to genes encoding Complex III, OCIAD1 (Ovarian Carcinoma Immunoreactive Antigen Domain containing-1) was identified as a strongly sensitizing hit. OCIAD1 encodes a poorly characterized predicted transmembrane protein (Figure 3A) that is aberrantly expressed in ovarian carcinomas and implicated in the regulation of mitochondrial metabolism via Complex I (Shetty et al., 2018). We validated the antimycin A-sensitizing phenotype of OCIAD1 by performing a growth competition assay in K562 cells using CRISPRi cell lines stably expressing an individual sgRNA against OCIAD1 (sgRNA#2). This sgRNA was identified in our screen and effectively silenced OCIAD1 expression (Figure 1C). Silencing OCIAD1 selectively compromised growth of antimycin A-treated cells, but not growth of oligomycin-treated cells (Figure 1D), suggesting, together with our screen data, that OCIAD1 knockdown specifically sensitizes cells to inhibition of Complex III.

### OCIAD1 is required for the assembly of Complex III

We assessed whether OCIAD1 regulates the assembly and/or stability of mitochondrial respiratory complexes using blue-native polyacrylamide gel electrophoresis (BN-PAGE), followed by Western blotting using antibodies directed against core constituents of respiratory Complexes I-V (Figure 2). Mitochondria were isolated from K562 control or OCIAD1 knockdown cells grown in galactose and respectively expressing a non-targeting sgRNA or sgRNA#2 against OCIAD1. We also analyzed mitochondria isolated from K562 OCIAD1 knockdown cells in which OCIAD1 expression had been reintroduced to near endogenous levels using lentiviral delivery (Figure 2A). There were no significant defects observed in the assembly of Complexes I, II, IV or V (Figure 2B, C, E and F). By contrast, we observed a selective defect in Complex III assembly in cells depleted of OCIAD1 (Figure 2D). The abundance of CIII_2_ was significantly reduced in mitochondria from OCIAD1 knockdown cells and restored to wildtype levels by OCIAD1 reintroduction, indicating that this defect is specific to loss of OCIAD1 function (Figure 2D, CIII_2_). At steady state, Complex III is an obligate dimer (CIII_2_) that participates with Complex I and IV (CI and CIV) to form higher-order assemblies in mitochondria, known as supercomplexes (CIII_2_CIV, CICIII_2_, CICIII_2_CIV). We also observed a smaller but significant reduction in CIII_2_ supercomplex assemblies (Figure 2D, SC). Mitochondrial respiratory chain complexes proteins are unusually long-lived (Fornasiero et al., 2018) and thus, the smaller impact of OCIAD1 on supercomplexes might be a consequence of enhanced stability of supercomplexes compared to individual complexes. In mitochondria depleted of OCIAD1, we also observed a reduction in a species whose mass/migration was consistent with the CIII_2_CIV supercomplex (Figure 2D, #). However, we did not detect a coincidental decrease in abundance of the co-migrating Complex IV species by Western blot analysis of COX4, a Complex IV marker (compare # and CIII_2_CIV in Figure 2D and 2E). The identity of this higher-order OCIAD1-sensitive Complex III species remains unknown.

**Figure 2.**
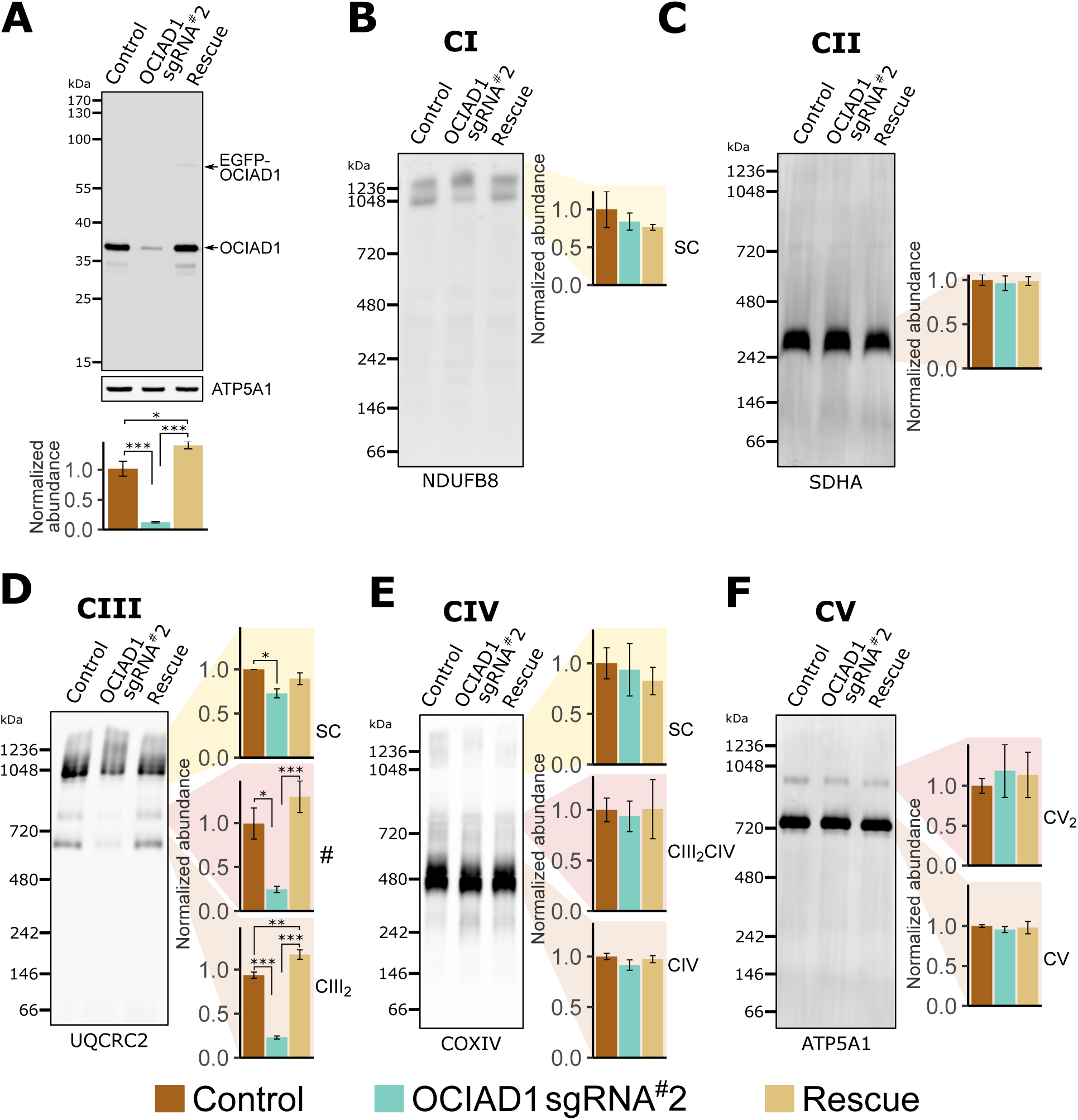
OCIAD1 is required for CIII_2_ assembly. A) Western blot showing CRISPRi silencing of OCIAD1 protein expression (12.47 ± 1.06% of control) in K562 cells. Rescue of OCIAD1 (141.20 ± 6.07% of control) by lentivirus transduction with a P2A multicistronic vector with high cleavage efficiency (98.89 ± 0.12%). The upper band (EGFP-OCIAD1) represents intact fusion gene product. ATP5A1 served as loading control. B-F) OCAID1 is selectively required for Complex III assembly. BN-PAGE analysis of digitonin-solubilized mitochondria followed by Western blotting using NDUFB8 (Complex I), SDHA (Complex II), UQCRC2 (Complex III), COXIV (Complex IV), and ATP5A1 (ATP synthase). The ATP5A1 signal from monomeric CV (F) was used as a loading control to quantify UQCRC2 intensities (D) as both proteins were probed on the same membrane. Values represent normalized intensity ± SEM (n = 3). Asterisks (*p < 0.05, **p < 0.01, or ***p < 0.001) correspond to the adjusted (FDR) p-values from the post-ANOVA pairwise t-test.

### OCIAD1 is a mitochondrial inner membrane protein

OCIAD1 is annotated as a mitochondrial protein by the MitoCarta 3.0 inventory (Rath et al., 2020), but also has been reported to localize to endosomes and peroxisomes (Antonicka et al., 2020; Mukhopadhyay et al., 2003; Sinha et al., 2018). Consistent with the MitoCarta repository, we found that OCIAD1 primarily localized to mitochondria in U2OS cells, as evidenced by indirect immunofluorescence analysis using validated polyclonal OCIAD1 antibodies (Figure 3B). Following extraction of peripheral membrane proteins with carbonate treatment of increasing pH, OCIAD1 and the known inner membrane protein TIM50 (Yamamoto et al., 2002) both remained in the membrane pellet fraction isolated by differential centrifugation (Figure 3C). In contrast, the peripheral membrane proteins ATP5A1 and SDHA were readily extracted and found in the supernatant (Figure 3C). These data indicate that OCIAD1 is an integral membrane protein, consistent with the presence of two predicted transmembrane domains (Figure 3A). Recently, OCIAD1 was suggested to reside in the outer mitochondrial membrane (OMM) based on proximity labeling (Antonicka et al., 2020; Lee et al., 2017). To further investigate the localization of OCIAD1, we conducted proteinase K protection assays on freshly isolated mitochondria. Whereas the validated OMM protein TOM70 was digested by treatment of intact mitochondria with proteinase K, OCIAD1 was resistant to degradation (Figure 3D). OCIAD1 was however susceptible to proteolytic degradation after compromising mitochondrial outer membrane integrity by hypo-osmotic treatment, similar to other inner mitochondrial membrane (IMM) proteins (Figure 3D). Overall, these results demonstrate that OCIAD1 is an integral IMM protein.

**Figure 3.**
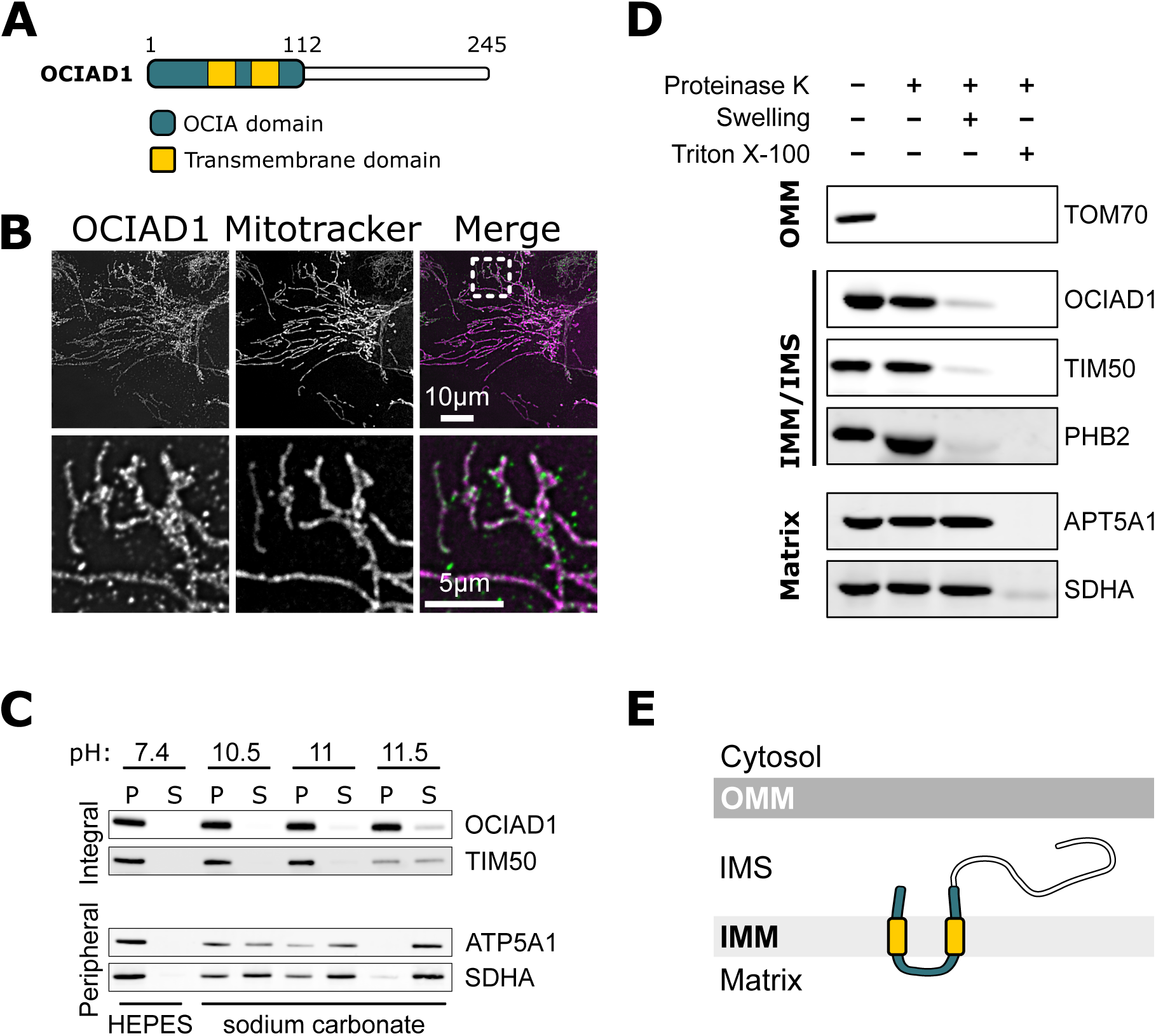
OCIAD1 is an inner mitochondrial membrane protein. A) Schematic illustration of OCIAD1domain organization. B) Representative images of fixed U2OS cells stained with Mitotracker (magenta) and immunolabeled using anti-OCIAD1 antibodies (green). Lower panel is a magnification of the inset shown in the upper panel. C) OCIAD1 is an integral membrane protein. Sodium carbonate extraction fractions (pH 10.5-11.5) immunoblotted with anti-OCIAD1, anti-TIM50, anti-ATP5A1, and anti-SDHA antibodies. P and S indicate pellet and soluble fractions, respectively. D) OCIAD1 localizes to the inner membrane. Protease protection assay fractions immunoblotted with anti-OCIAD1, anti-prohibitin 2 (PHB2), anti-TIM50, anti-ATP5A1, and anti-SDHA antibodies. (OMM: outer mitochondrial membrane, IMM: inner mitochondrial membrane, IMS: intermembrane space). E) Schematic illustration of OCIAD1 topology within the inner membrane.

Next, we sought to determine the topology of the OCIAD1 protein within the IMM using proteinase K as the amino acid sequence between the two predicted OCIAD1 transmembrane domains contains predicted proteinase K cleavage sites. Analysis of OCIAD1 deletion constructs by Western blotting analysis using polyclonal anti-OCIAD1 antibodies identified the last 25 amino acids of the OCIAD1 C-terminus as the antigenic determinant (Figure 3-figure supplement 1A,B). OCIAD1 proteolytic fragments were not observed by Western analysis of mitoplasts treated with proteinase K (Figure 3-figure supplement 1C). Thus, given the location of the OCIAD1 epitope, this observation suggests that the C-terminus was degraded and therefore localized to the intermembrane space. We also tested OCIAD1 localization and topology using a bipartite split GFP complementation assay, in which the 11 stranded β-barrel GFP fluorophore is reconstituted from a separately expressed N-terminal β-strands (GFP_1-10_) and a C-terminal 16 amino acid β-strand (GFP_11_), as previously described (Hyun et al., 2015). Specifically, we created U2OS cells stably expressing GFP_1-10_ targeted either to the matrix or intermembrane space (IMS) and transiently expressed proteins tagged with GFP_11_. We validated this system by expressing known matrix and IMS proteins, CoQ9 and MICU1, respectively, with C-terminal GFP_11_ tags and measuring the efficiency of GFP complementation by flow cytometry (Figure 3-figure supplement 1D). CoQ9-GFP_11_ complemented GFP only when expressed in matrix-targeted GFP_1-10_ cells, consistent with its matrix localization (Johnson et al., 2005). Conversely, MICU1-GFP_11_ only produced GFP signal when expressed in IMS-targeted GFP_1-10_ cells, consistent with its localization to the IMS (Hung et al., 2014; Tsai et al., 2016). With a validated topology assay in hand, we transiently expressed OCIAD1 tagged with GFP_11_ at either the N- or C-terminus and observed GFP fluorescence signal only in the IMS-targeted GFP_1-10_ cells (Figure 3-figure supplement 1D). Together, these data indicate that OCIAD1 is a transmembrane protein embedded in the mitochondrial inner membrane with its N- and C-termini facing the IMS (Figure 3E).

### OCIAD1 interacts with the supramolecular prohibitin complex

To gain insight into how OCIAD1 facilitates steady state Complex III assembly, we mapped its interactome using affinity enrichment mass spectrometry (AE-MS). We immunopurified OCIAD1 from DSP-crosslinked cell lysates prepared from K562 OCIAD1 knockdown cells and K562 OCIAD1 knockdown cells rescued with wildtype OCIAD1 and analyzed the eluates by label-free quantitative mass spectrometry (Figure 4A-source data 3). We identified Complex III subunits and assembly factors, which supports our BN-PAGE data indicating that OCIAD1 regulates Complex III assembly (Figure 4A, in green). In addition, we identified subunits of the prohibitin complex, PHB1 and PHB2, as potential OCIAD1 interactors (Figure 4A, in dark purple), consistent with previously published work (Richter-Dennerlein et al., 2014). We also identified several prohibitin interactors, including C1QBP, COX4I1, and DNAJC19, the mitochondrial m-AAA proteases AFG3L2 and SPG7, as well as the AFG3L2-interactor MAIP1, and the protease IMMP2L, all previously identified in published studies examining the prohibitin interactome (Richter-Dennerlein et al., 2014; Yoshinaka et al., 2019) (Figure 4A, in light green). Prohibitins form large hetero-oligomeric ring complexes composed of assemblies of PHB1/PHB2 dimers in the inner membrane of mitochondria (Tatsuta et al., 2004). These complexes are thought to constitute molecular scaffolds that define functional domains to regulate the lateral distribution of membrane lipids and proteins within the inner mitochondrial membrane (Osman et al., 2009; Richter-Dennerlein et al., 2014). Prohibitin structures associate with the inner membrane matrix-AAA protease to modulate their activity in both the specific processing of inner membrane proteins and the targeted degradation of unassembled inner membrane proteins (Bonn et al., 2011; Ehses et al., 2009; Koppen et al., 2009; Li et al., 2019; Merkwirth et al., 2008; Steglich et al., 1999).

**Figure 4.**
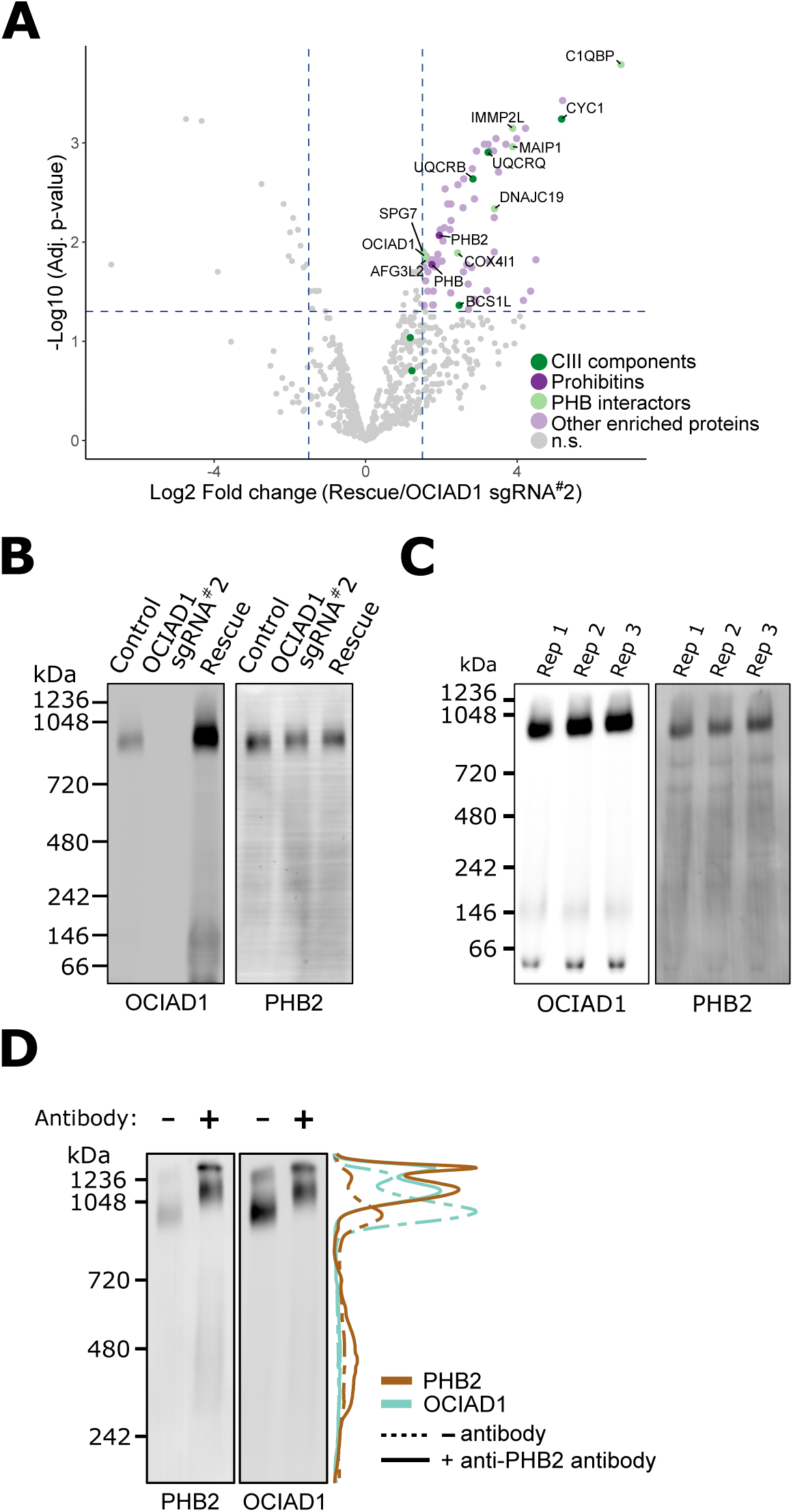
OCIAD1 forms a complex with prohibitin supramolecular assemblies. A) Volcano plot showing the statistical significance (-log_10_ FDR adjusted p-value; y axis) vs log_2_ fold change (x axis) of proteins enriched in OCIAD1 pull-down performed on DSP-crosslinked K562 cell lysates from OCIAD1 knockdown cells and OCIAD1 knockdown cells rescued with wildtype OCIAD1. Proteins with a log_2_ fold change ≥ 1.5 and an adjusted p-value < 0.05 were considered significantly enriched. (n = 3, n.s. = non-significantly enriched). B) BN-PAGE of LMNG detergent-solubilized mitochondrial membranes isolated from U2OS control, OCIAD1 knockdown, and OCIAD1 knockdown cells rescued with wildtype OCIAD1. The membrane was immunoblotted with anti-OCIAD1 and anti-prohibitin 2 antibodies. C) BN-PAGE of LMNG detergent-solubilized mitochondrial membranes isolated from U2OS control cells (n = 3) and immunoblotted with anti-OCIAD1 and anti-prohibitin 2 antibodies. Electrophoresis was stopped before elution of the migration front to calculate the fraction of OCIAD1 that associates with PHB2 assemblies (66.91 ± 0.35%). D) Mitochondria from K562 cells solubilized with LMNG and pre-incubated with anti-Phb2 antibodies (solid line) or vehicle (dotted line) were analyzed by BN-PAGE and immunoblotted with anti-OCIAD1 and anti-prohibitin 2 antibodies. Line scan traces represent the distribution profile of Phb2 (brown) and OCIAD1 (light blue).

We also examined the molecular features of native OCIAD1 by BN-PAGE to assess potential interactors. Specifically, LMNG-solubilized mitochondria isolated from K562 control and OCIAD1 knockdown cells, as well as K562 rescue cells expressing wildtype OCIAD1, were analyzed by BN-PAGE followed by Western analysis using anti-OCIAD1 antibodies (Figure 4B, left panel). This analysis demonstrated that the majority of OCIAD1 associates with large 0.9-1MDa supramolecular assemblies that co-migrate with prohibitin (Figure 4B,C). To test whether OCIAD1 and prohibitin interact within the 0.9-1MDa supramolecular assemblies, we performed an in-gel mobility shift assay. For this purpose, LMNG-solubilized mitochondrial membranes from K562 cells were incubated with either vehicle alone or with anti-PHB2 antibody and subjected to BN-PAGE, followed by Western analysis using anti-PHB2 or OCIAD1 antibodies. Pre-incubation of mitochondrial membranes with anti-PHB2 antibody, but not vehicle, retarded the migration of both PHB2 and OCIAD1 supramolecular assemblies to a similar degree (Figure 4D, compare dashed and solid lane line scans, respectively). Thus, together our data indicate that OCIAD1 associates with prohibitin complexes in the inner mitochondrial membrane.

### The OCIAD1 paralog, OCIAD2, has similar topology and interacts with prohibitin but is not functionally redundant with OCIAD1

OCIAD1 has a paralog in vertebrates, OCIAD2, which likely arose from tandem gene duplication of a common ancestor around 435–500 million years ago (Sinha et al., 2018). The paralogs share domain structure and significant homology (Figure 4-figure supplement 1A) and have been reported to hetero-oligomerize (Sinha et al., 2018), suggesting a shared function. Using indirect immunofluorescence, carbonate extraction, and protease protection analysis, we showed that, as expected, OCIAD2 localized to mitochondria with a topology similar to OCIAD1 (Figure 4-figure supplement 1B-D).

We examined whether OCIAD2, like OCIAD1, functions in Complex III assembly. OCIAD2 was not a hit in our antimycin A screen (Figure 4-figure supplement 2A) and Western blot analysis indicated that the K562 cells used in our CRISPRi screen do not express OCIAD2 (Figure 4-figure supplement 2B). Therefore, we used U2OS cells, which express both paralogs, and generated individual and double OCIAD1/OCIAD2 knockdown cell lines, by identifying OCIAD2 shRNAs that efficiently silenced OCIAD2 expression (Figure 4-figure supplement 2C, shRNA1-3). Similar to our results in K562 cells, expression of sgRNA#2 against OCIAD1 effectively suppressed OCIAD1 expression in U2OS cells (0.86 ± 0.43% of control) but did not affect OCIAD2 expression (104.38 ± 18.48% of control; Figure 4-figure supplement 2D). Conversely, expressing a shRNA targeting OCIAD2 selectively silenced OCIAD2 expression in U2OS cells (8.17 ± 2.75% of control), but did not alter OCIAD1 levels (92.89 ± 8.46% of control; Figure 4-figure supplement 2D). We next examined the abundance of CIII_2_ in the different cell lines by BN-PAGE analysis of mitochondria isolated from cells grown in glucose-free media containing galactose. Knockdown of OCIAD1 in U2OS cells decreased steady-state levels of CIII_2_ relative to control cells, consistent with our observations in K562 cells. In contrast, knockdown of OCIAD2 did not affect CIII_2_ levels (Figure 4-figure supplement 2E). Given that this analysis was performed on cells grown in glucose-free media containing galactose, we considered whether the role of OCIAD1 and OCIAD2 in CIII_2_ assembly was modulated by carbon source/metabolism. We used BN-PAGE to monitor CIII_2_ levels in U2OS cells grown in media containing glucose (Figure 4-figure supplement 3). Similar to galactose media, CIII_2_ abundance was markedly reduced in mitochondria from OCIAD1 knockdown cells grown in glucose media, but not in OCIAD2 knockdown cells, indicating that OCIAD1, but not OCIAD2, affects the assembly of Complex III under our experimental conditions.

To gain insight into OCIAD1 and OCIAD2 function, we also used untargeted quantitative mass spectrometry to compare the whole-cell proteomes of control U2OS cells, U2OS cells with individual or double OCIAD1/OCIAD2 knockdown, and OCIAD1 knockdown U2OS cells in which OCIAD1 expression was reintroduced by lentiviral delivery. Overall, the proteome was resilient to loss of OCIAD1 and OCIAD2 expression, as only 38 proteins were significantly affected in at least one of the different cell lines (Figure 4-figure supplement 4A). As expected, OCIAD1 and OCIAD2 were significantly downregulated in the individual and double knockdown cell lines, while GFP was only observed in OCIAD1 knockdown cell line in which OCIAD1 expression was reintroduced by lentiviral transduction using GFP as a selection marker. Hierarchical clustering of significantly affected proteins identified a small cluster tightly associated with OCIAD1 (Figure 4-figure supplement 4A, red box), containing 4 of the 10 subunits of Complex III, including UQCRC1, UQCRC2, UQCRB, and cytochrome *c_1_* (CYC1), as well as COX7A2L, which regulates Complex III biogenesis by promoting the assembly of CIII_2_ with CIV to form the CIII_2_CIV supercomplex (Lobo-Jarne et al., 2018). All proteins in this cluster were selectively downregulated in the individual OCIAD1 knockdown and OCIAD1/OCIAD2 double knockdown cell lines, but unaffected in the OCIAD1 rescued cell line and in OCIAD2 knockdown cells (Figure 4-figure supplement 4A). We validated these observations using Western blotting and showed that steady-state levels of UQCRC1, UQCRC2, and CYC1 were reduced in mitochondria isolated from OCIAD1 knockdown cells, but not OCIAD2 knockdown cells (Figure 4-figure supplement 4B and Figure 5 E,F).

**Figure 5.**
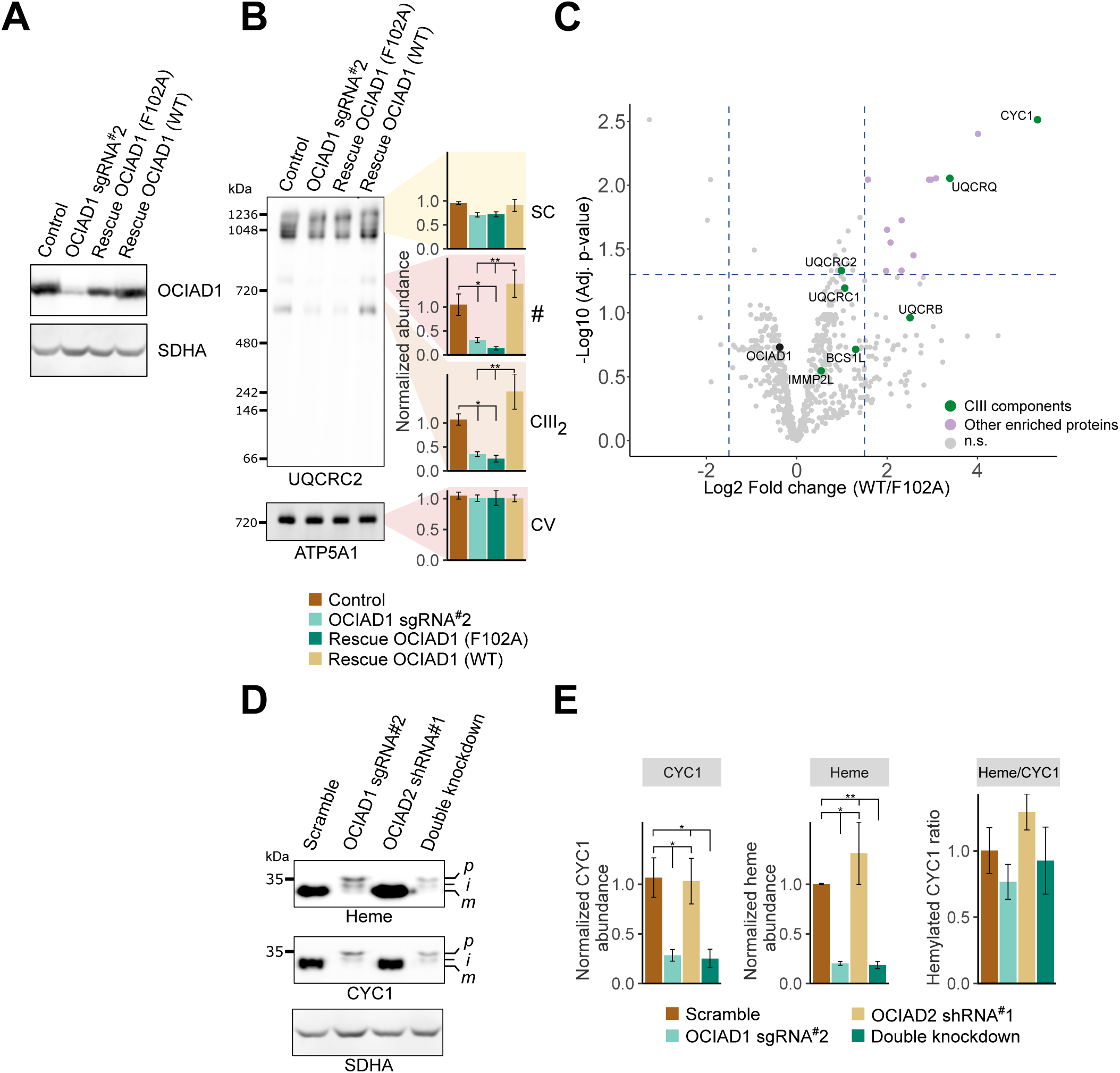
OCIAD1 regulates the maturation of cytochrome *c*_1_. A) Western blot showing OCIAD1 expression levels in K562 OCIAD1 knockdown cells rescued with either wildtype OCIAD1 or mutant (F102A) OCIAD1. B) Blue-native PAGE analysis showing that the F102A point mutant fails to rescue the CIII_2_ assembly defect. C) Volcano plot showing proteins enriched in OCIAD1 pull-down performed in DSP-crosslinked cell lysate from K562 OCIAD1 knockdown cells rescued with either wildtype or F102A OCIAD1. (n.s = non-significantly enriched) D) Western blot analysis of U2OS mitochondrial membranes solubilized in digitonin. Heme was detected by chemiluminescence before immunoblotting the membrane with the indicated antibodies. E) Quantification of CYC1 (left) and heme (middle) levels from blot shown in figure 5E. Right panel shows the proportion of CYC1 that is hemylated. Values represent normalized intensity ± SEM (n = 3). Asterisks (*p < 0.05, **p < 0.01, or ***p < 0.001) correspond to the adjusted (FDR) p-values from the post-ANOVA pairwise t-test.

Although OCIAD2 does not have a measurable effect on Complex III biogenesis under our conditions, BN-PAGE analysis and Western analysis of mitochondria from U2OS cells using anti-OCIAD2 antibody demonstrated that, like OCIAD1, OCIAD2 co-migrates with prohibitin complexes independent of OCIAD1 (Figure 4-figure supplement 1E). Reciprocally, OCIAD2 is not required for the association of OCIAD1 with prohibitin complexes as OCIAD1 migrated as 0.9-1MDa assemblies in K562 cells that do not express OCIAD2 (Figure 4C). Therefore, our data suggest that OCIAD1 and OCIAD2 paralogs experienced functional diversification during evolution. However, as OCIAD1 and OCIAD2 have been reported to interact (Sinha et al., 2018), we cannot exclude the possibility that OCIAD2 may modulate the role of OCIAD1 in Complex III regulation in a context-dependent manner.

### OCIAD1 is required for the processing of cytochrome c_1_

To determine the functional significance of OCIAD1 interactors, we asked which interactions were modulated by OCIAD1 loss-of-function in Complex III assembly. To identify OCIAD1 loss-of-function alleles, we performed a structure-function analysis by initially creating tiled deletions along the C-terminus of OCIAD1 and identified a segment of the conserved OCIA domain essential for OCIAD1 function in CIII_2_ assembly (Figure 4-figure supplement 5A,B). Sequence alignments of OCIAD1 genes from distant phylogenetic species identified a highly conserved phenylalanine residue (F102) within this region (Figure 4-figure supplement 5C, red box), which we mutated to an alanine. Western blot analysis of mitochondria isolated from K562 OCIAD1 knockdown cells exogenously expressing either wildtype (WT) or mutated (F102A) OCIAD1 indicated that OCIAD1 F102A was expressed at near endogenous levels in rescued cells (Figure 5A). We examined Complex III assembly by BN-PAGE analysis of mitochondria isolated from K562 OCIAD1 knockdown cells rescued with either wildtype or OCIAD1 F102A. In contrast to cells expressing wildtype OCIAD1, cells expressing the F102A mutant had decreased levels of Complex III, indicating that F102A constitutes a loss-of-function mutation (Figure 5B).

We compared the interactomes of wildtype OCIAD1 and OCIAD1 F102A using AE-MS. OCIAD1 was immunopurified from DSP-crosslinked whole-cell lysates prepared from OCIAD1 knockdown cells rescued with either wildtype OCIAD1 or OCIAD1 F102A and analyzed by label-free quantitative mass spectrometry, as in Figure 4A. This analysis revealed a selective enrichment of CYC1, one of three catalytic CIII_2_ subunits, in cells rescued with wildtype OCIAD1 versus OCIAD1 F102A (Figure 5C-source data 4). These data suggest that the function of OCIAD1 in Complex III assembly is dependent on its interaction with CYC1.

CYC1 contains a single covalently attached heme prosthetic group that facilitates the transfer of electrons from the Rieske iron–sulfur protein to cytochrome *c*. It is synthesized on cytosolic ribosomes as an apoenzyme precursor with a bipartite signal sequence that is processed in two steps during import. The CYC1 precursor is first processed to an intermediate form by the matrix metalloprotease (MPP), which removes the matrix targeting sequence (Gasser et al., 1982; Ndi et al., 2018; Nicholson et al., 1989). The matrix targeting sequence is followed by a stretch of hydrophobic residues that functions as a signal that stops the translocation of the mature protein across the inner membrane, allowing CYC1 to localize to the intermembrane space. The stop transfer signal is processed by IMMP2L, a signal peptidase-like protease (Gasser et al., 1982; Ndi et al., 2018; Nicholson et al., 1989; Nunnari et al., 1993). IMMP2L processing requires the covalent addition of a heme moiety to CYC1, catalyzed by holocytochrome c-type synthase (HCCS), and completes the formation of mature holo-CYC1 (Ndi et al., 2018; Nicholson et al., 1989).

IMMP2L was identified in our OCIAD1 interactome analysis (Figure 4A) and also in a prohibitin interactome analysis (Yoshinaka et al., 2019). Therefore, we assessed whether OCIAD1 regulates CYC1 maturation using Western blotting analysis of digitonin-solubilized mitochondria. CYC1 levels were significantly reduced in OCIAD1 knockdown cells (Figure 5 D and E), consistent with our unbiased mass spectrometry data (Figure 4-figure supplement 4A). In addition, two larger molecular weight CYC1 species, likely corresponding to the precursor and intermediate forms, accumulated in OCIAD1 knockdown cells as compared to control cells (Figure 5 D and E). This phenotype was also observed in OCIAD1 knockdown cells expressing the OCIAD1 loss-of-function truncation (Δ97-115), which failed to rescue CIII_2_ assembly (Figure 5-figure supplement 1A). These data indicate that OCIAD1 function is required for normal CYC1 processing. Given that hemylation is required for IMMP2L processing of CYC1 (Nicholson et al., 1989), we also examined whether CYC1 hemylation was dependent on OCIAD1 function. As CYC1 contains a covalently linked heme moiety, we directly assessed CYC1 hemylation by chemiluminescence of mitochondrial fractions analyzed by SDS-PAGE as previously described (Dorward, 1993; Feissner et al., 2003). The slower migrating CYC1 species that accumulate in OCIAD1 knockdown cells and OCIAD1 Δ97-115 truncated cells were fully hemylated (Figure 5 D,E-figure supplement 1A). The hemylation levels of the slower-migrating CYC1 species in OCIAD1 knockdown cells were proportional to CYC1 abundance, indicating that OCIAD1 knockdown cells have a CYC1 hemylation ratio comparable to mature CYC1 in control cells (Figure 5E). These data indicate that OCIAD1 is not required for CYC1 hemylation. The CYC1 maturation defect was not detected in OCIAD2 knockdown cells and, thus, was specific to loss of OCIAD1 function (Figure 5D and E). This is consistent with our results showing that OCIAD2 was not required for CIII_2_ assembly and the conclusion that OCIAD1 and OCIAD2 are functionally divergent (Figure 4-figure supplement 2 and 3).

To further investigate the role of OCIAD1 in CYC1 processing, we examined the status of CYC1 in CIII_2_ in OCIAD1 knockdown cells. We measured the hemylation efficiency of CYC1 in CIII_2_ resolved by native PAGE (Figure 5-figure supplement 1B). Although CIII_2_ levels were reduced in OCIAD1 knockdown cells, the extent of hemylation in CIII_2_ was comparable to that of wildtype cells (compare Figure 5F and Figure 5-figure supplement 1C). We also used 2D-native/SDS-PAGE and found that CIII_2_ from OCIAD1 knockdown cells contains higher molecular weight CYC1 species (Figure 6A). Thus, unprocessed but hemylated CYC1 is incorporated in CIII_2_ in OCIAD1 knockdown cells.

**Figure 6.**
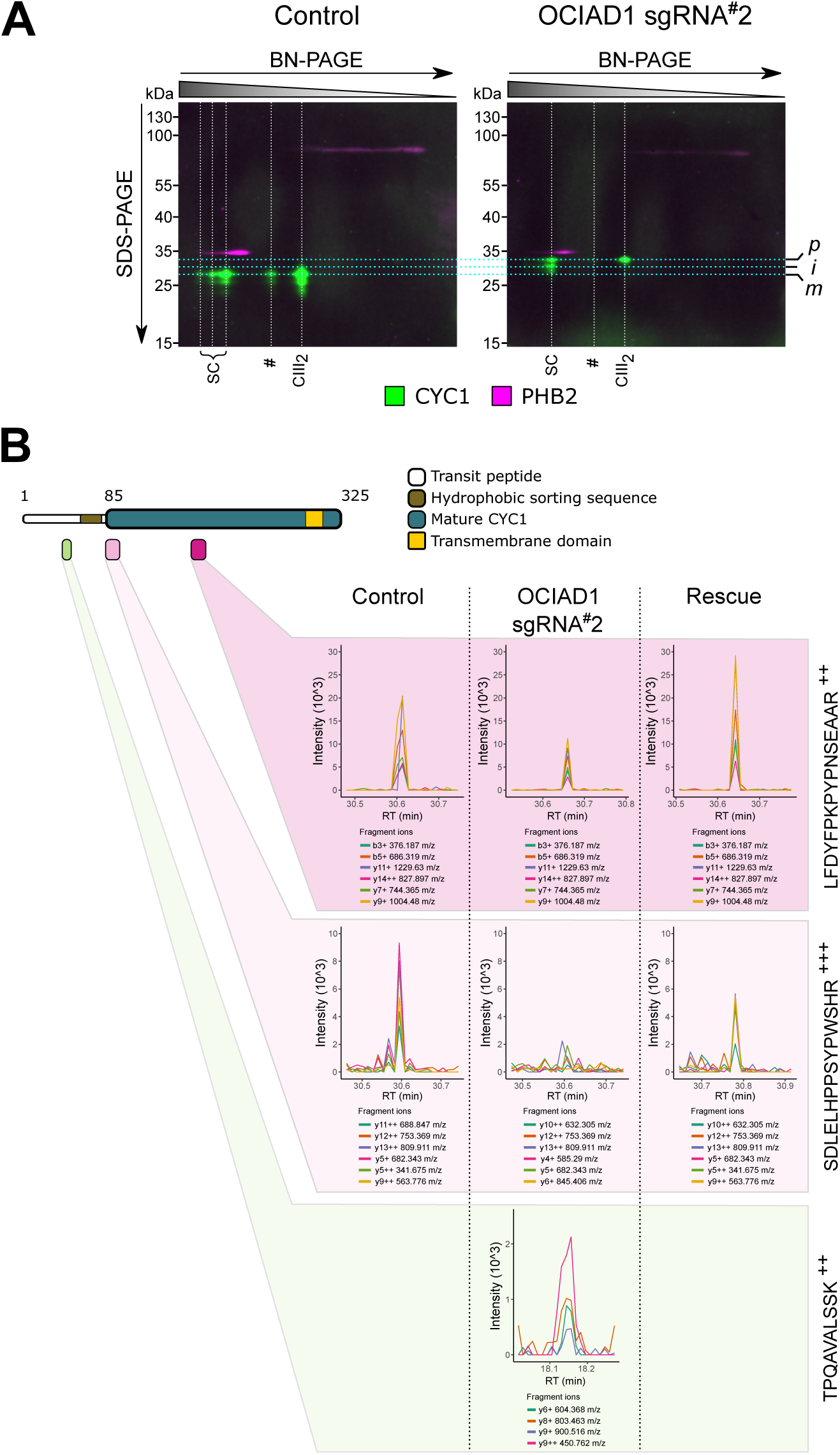
OCIAD1 regulates IMMP2L-dependent proteolytic processing of cytochrome *c_1_*. A) 2D-native/SDS-PAGE analysis of mitochondrial membranes isolated from K562 control and OCIAD1 knockdown cells and immunoblotted with CYC1 and PHB2 antibodies. CIII_2_ assemblies from OCIAD1 knockdown cells contained immature CYC1 of higher molecular weight. PHB2 staining served as an internal molecular size reference. Light blue horizontal lines represent the size of putative precursor (*p*), intermediate (*i*), and mature (*m*) CYC1. White vertical lines represent the different high-order CIII_2_ assemblies. B) Extracted MS2 fragment ion chromatograms (XIC) for three diagnostic CYC1 peptides detected by diaPASEF mass spectrometry in BN-PAGE gel slices excised from control cells, OCIAD1 knockdown cells, and OCIAD1 knockdown cells rescued with wildtype OCIAD1. Individual peptides displayed highly correlated fragment ion co-elution profiles strongly supportive of peptide identification. The TPQAVALSSK^++^ peptide (bottom panel), located at the N-terminus of the CYC1 hydrophobic sorting sequence, was only identified in CIII_2_ assemblies from OCIAD1 knockdown cells. Conversely, the SDLELHPPSYPWSHR^+++^ peptide (middle panel), which uniquely identifies the N-terminus of mature CYC1 but is not present in the tryptic digest of the CYC1 precursor, was reliably detected in CIII_2_ assemblies from control and OCIAD1 knockdown cells rescued with wildtype OCIAD1, but not from OCIAD1 knockdown cells. An internal peptide (LFDYFPKPYPNSEAAR^+++^, top panel) common to all CYC1 species (precursor, intermediate, mature) was detected in all cell lines, albeit at lower levels in OCIAD1 cells as expected.

To better characterize the nature of the CYC1 processivity defect in CIII_2_, we identified CYC1 peptides using mass spectrometry analysis of BN-PAGE gel slices containing CIII_2_ assemblies ranging from ∼600-900kDa excised from control cells, OCIAD1 knockdown cells, and OCIAD1 knockdown cells rescued with wildtype OCIAD1 (Figure 6B). An internal peptide from mature CYC1 (LFDYFPKPYPNSEAAR) was reliably identified in CIII_2_ assemblies from mitochondria isolated from all cell types, albeit at lower levels in knockdown cells. This is consistent with our whole-cell proteomics and Western blot results showing reduced steady-state levels of CYC1 in OCIAD1 knockdown cells and an overall reduction in CIII_2_ assemblies by BN-PAGE. We also identified a peptide (TPQAVALSSK), N-terminal to the CYC1 hydrophobic bipartite sequence, that was uniquely detected in CIII_2_ assemblies isolated from OCIAD1 knockdown cells (Figure 6B, lower panel), consistent with the accumulation of the precursor form of CYC1 in OCIAD1 knockdown cells. Conversely, a peptide (SDLELHPPSYPWSHR) representing the N-terminus of mature CYC1, as determined by N-terminone sequencing analysis of the human mitochondrial proteome (Vaca Jacome et al., 2015), was only reliably identified in CIII_2_ assemblies from control and rescued cells, but not from OCIAD1 knockdown cells (Figure 6B, middle panel). This peptide is not preceded by an arginine or lysine residue and thus was not produced by tryptic digestion of the CYC1 precursor. Therefore, this peptide distinctly identifies the N-terminus of the mature version of CYC1 (Vaca Jacome et al., 2015). Taken together, our results are consistent with OCIAD1 regulating the proteolytic processing and maturation of the holocytochrome *c_1_* precursor.

## Discussion

Our data indicate that OCIAD1 is a conserved regulatory determinant of CIII_2_ assembly that controls the proteolytic processing of holocytochrome *c_1_*. Cytochrome bc1 complexes are highly conserved, found in photosynthetic and respiring bacterial plasma membranes of phylogenetically distant species, as well as in eukaryotic cells in mitochondria and in chloroplasts as the related cytochrome b6f complex (Trumpower, 1990). The comparison of high resolution atomic models of cytochrome bc1 complexes in plants, fungi, and mammals revealed that despite their modest sequence homology, they exist as dimers (CIII_2_) displaying exceptional structural conservation of all three catalytic subunits (Maldonado et al., 2021). In all CIII_2_ atomic models, mature holocytochrome *c_1_* possesses one C-terminal transmembrane helix and an N-terminal domain composed of 6 α-helices and 2-strand β-sheet extending in the IMS (Maldonado et al., 2021; Xia et al., 1997). This topology is achieved via the highly conserved process of CYC1 maturation. CYC1 contains a bipartite targeting signal composed of two sequential N-terminal presequences: a mitochondrial targeting signal (MTS) processed in the matrix by the MPP and a hydrophobic inner membrane sorting domain. Following hemylation of CYC1 by the heme lyase HCCS, the hydrophobic inner membrane sorting domain is processed by IMMP2L, a subunit of the inner membrane signal peptidase complex (Arnold et al., 1998; Nicholson et al., 1989; Nunnari et al., 1993; Römisch et al., 1987; Sadler et al., 1984; van Loon et al., 1987; Wachter et al., 1992). The MTS of CYC1 is required for its targeting to mitochondria; however, its removal by MPP is not required for the heme-dependent maturation of CYC1 by IMMP2L. Thus, both precursor and intermediate forms of CYC1 can be hemylated and the bipartite sequence can be cleaved in a single step by the IMMP2L, without the removal of the MTS by the MPP (Nicholson et al., 1989). Consistent with this, two fully hemylated CYC1 species of higher molecular weight accumulate in OCIAD1 knockdown and mutant cells and likely represent the precursor and intermediate holocytochrome *c_1_*. Our OCIAD1 interactome analysis identified IMMP2L (Figure 4A), which is required for the second processing step of CYC1. IMMP2L was also identified in a prohibitin interactome analysis (Yoshinaka et al., 2019). Thus, overall, our results suggest that OCIAD1 regulates CYC1 processing by both MPP and IMMP2L.

OCIAD1 and its related paralog OCIAD2 are highly conserved in metazoans. In addition, remote protein homology detection using HHpred analysis found homology similarity between OCIAD1 and the yeast protein COX20. COX20 has a similar size and topology to OCIAD1. Both proteins are integral inner membrane proteins with N- and C-termini located in the intermembrane space (Tzagoloff et al., 2000). COX20 is a chaperone involved in the biogenesis of CIV where it binds to newly synthesized mitochondrially-encoded Cox2. COX20 helps present Cox2, a CIV subunit synthesized as a precursor protein in yeast, to the inner membrane peptidase complex to facilitate the proteolytic removal of its N-terminal presequence by Imp2 (Elliott et al., 2012; Nunnari et al., 1993; Tzagoloff et al., 2000). Our study indicates that OCIAD1 serves a conserved function by facilitating the proteolytic processing of CYC1.

Processing of CYC1 to its mature form is not essential for its function in CIII_2_ as mitochondrial respiration, including CIII_2_ activity, is not affected in IMMP2L mutant mice (Bharadwaj et al., 2014; Lu et al., 2008). Consistent with this, we found that immature holocytochrome *c_1_* can be successfully incorporated into CIII_2_ and CIII_2_-containing supercomplexes in OCIAD1 knockdown and mutant cells. In this context, the functional significance of proteolytic processing of cytochrome *c_1_* is unclear, although it is an evolutionary conserved process. It is possible that CIII_2_ complexes containing immature holocytochrome *c_1_* have increased superoxide production, which can be detrimental to mitochondrial function (Lu et al., 2008). However, in contrast to IMMP2L mutant mice (Bharadwaj et al., 2014; Lu et al., 2008), steady-state levels of CYC1 were also substantially reduced in OCIAD1 knockdown cells, consistent with the observed antimycin A-sensitization growth phenotype of OCIAD1 knockdown cells. This observation suggests that OCIAD1 may function also as a chaperone that stabilizes newly imported CYC1. We demonstrate that under native conditions, a majority of OCIAD1 associates with prohibitins to form supramolecular complexes of ∼1MDa, consistent with published prohibitin interactomes (Richter-Dennerlein et al., 2014; Yoshinaka et al., 2019). OCIAD1 is likely a direct prohibitin interactor given that residue-to-residue contacts were identified between prohibitin and OCIAD1 peptides using cross-linking mass spectrometry analysis (Liu et al., 2018; Yoshinaka et al., 2019). Prohibitins are members of the SPFH superfamily of scaffold proteins and form large ring-like structures in membranes, which are thought to create functionally specialized protein and lipid domains within the crowded environment of the IMM (Osman et al., 2009). Consistent with this model, prohibitins, and the related SPFH scaffold, SLP2, have been shown to sequester inner membrane associated proteases to gate their access to proteolytic substrates (Merkwirth et al., 2008; Steglich et al., 1999). IMMP2L was identified in the prohibitin interactome by proximity labeling (Yoshinaka et al., 2019). Thus, we propose that within prohibitin assemblies OCIAD1 targets precursor CYC1 to the IMMP2L peptidase.

## Methods

### Cell culture

K562 cells and derivatives were cultured in “RPMI+glucose” (RPMI 1640 from HyClone (cat# SH30255F) or Gibco (cat# 72400047) supplemented with 10% fetal bovine serum (FBS), 100 units/mL penicillin, and 100 µg/mL streptomycin) or glucose-free “RPMI+galactose” (RPMI 1640 from Gibco (cat# 11879020) supplemented with 10mM galactose, 25mM HEPES, 10% FBS, 100 units/mL penicillin, 100 µg/mL streptomycin) where indicated. U2OS cells and derivatives, as well as HEK293T cells, were cultured in “DMEM+glucose” (DMEM from Gibco (cat# 12430054) supplemented with 10% FBS, 100 units/mL penicillin, 100 µg/mL streptomycin) or glucose-free “DMEM+galactose” (DMEM from Gibco (cat# 11966025) supplemented with 10mM galactose, 10% FBS, 100 units/mL penicillin, 100 µg/mL streptomycin) where indicated.

### Cloning and plasmid construction

Sequences of oligonucleotides used for cloning are provided in Supplementary table 1. Cloning was performed using Phusion or Platinum SuperFi high fidelity DNA polymerases (Thermo Scientific, cat# F530S and 12351010) and Gibson assembly master mix (New England BioLabs, cat# E2611). Individual OCIAD1 sgRNA and OCIAD2 shRNA vectors were generated by annealed oligo cloning of top and bottom oligonucleotides (Integrated DNA Technologies, Coralville, IA) into an optimized lentiviral pU6-sgRNA Ef1α-Puro-T2A-BFP vector digested with BstXI/BlpI (Addgene, cat# 84832) and a pLKO.1 backbone digested with AgeI/EcoRI (Addgene, cat# 26655), respectively. OCIAD1 was initially cloned from human cDNA into a pAcGFP-N1 vector (Clontech, Mountain View, CA). The GFP_1-10_ vectors were cloned by Gibson assembly into a FUGW lentiviral backbone (Addgene, cat# 14883) digested with BamHI/EcoRI. The MTS- and IMS-targeting sequences were ordered as gene blocks (Integrated DNA Technologies, Coralville, IA) and the GFP_1-10_ fragment was cloned from a pCMV-mGFP_1–10_ plasmid (Van Engelenburg and Palmer, 2010). The mitochondrial targeting signal (MTS) from yeast COX4 (a.a. 1-21) (Friedman et al., 2011) and the IMS-targeting signal from MICU1 (a.a. 1-60) (Gottschalk et al., 2019; Hung et al., 2014; Tsai et al., 2016) were chosen to target GFP_1-10_ to the matrix or IMS, respectively. The pGFP_11_-N1 and pGFP_11_-C1 vectors were cloned by Gibson assembly into a pEGFP-N1 backbone (Clontech, Mountain View, CA) digested with BamHI/NotI to replace the GFP gene with the GFP_11_ β-barrel. The GFP_11_ fragments were ordered as gBlocks (Integrated DNA Technologies, Coralville, IA) and contained a strategically located BamHI cloning site for easy N- or C-terminal tagging. CoQ9 and MICU1 genes were cloned from human cDNA and inserted into a BamHI-digested pGFP_11_-N1 vector by Gibson assembly to generate the pCoQ9-GFP_11_ and pMICU1-GFP_11_ plasmids. Similarly, OCIAD1 was amplified from the pAcGFP-OCIAD1 plasmid and cloned into BamHI-digested pGFP_11_-N1 and pGFP_11_- C1 vectors to create the pOCIAD1-GFP_11_ and pGFP_11_-OCIAD1 plasmids, respectively. OCIAD1 was also amplified from the pAcGFP-OCIAD1 plasmid and cloned into a XbaI/BamHI-digested pUltra-EGFP backbone (Addgene, cat# 24129) to generate a lentiviral vector expressing the GFP-OCIAD1 fusion gene containing a “self-cleaving” P2A sequence. The OCIAD1 F102A point mutant was generated from this pUltra-OCIAD1 vector using site-directed mutagenesis. We also generated an OCIAD1 construct with a C-terminal StrepII tag preceded by a TEV cleavage site. For this, OCIAD1 was amplified from the pAcGFP-OCIAD1 plasmid and inserted into a XbaI/BamHI-digested pUltra-EGFP vector by Gibson assembly, together with a gBlock (Integrated DNA Technologies, Coralville, IA) encoding the TEV-StrepII sequence. The OCIAD1 truncation constructs were generated by inverse PCR using the pAcGFP-OCIAD1 and pUltra-OCIAD1-TEV-StrepII vectors as templates. Finally, to generate lentiviral vectors expressing the truncated OCIAD1 isoforms with a C-terminal GFP tag, the entire OCIAD1-GFP cassette containing the deletion was amplified from the various pAcGFP-OCIAD1 truncated constructs and cloned by Gibson assembly into a FUGW plasmid digested with BamHI/EcoRI to remove its GFP gene.

### Lentivirus production, infection, and generation of cell lines

Lentivirus were generated by transfecting HEK293T cells with standard packaging vectors using TransIT-LT1 Transfection Reagent (Mirus Bio, Madison, WI) or Lipofectamine 2000 (Invitrogen, Carlsbad, CA) according to the manufacturer’s instructions. Briefly, HEK293T were plated in a 6 wells plate on day 0 (0.5×10^6^ cells per well) and transfected on day 1 with a liposome/DNA mixture containing the following packaging plasmids (0.1µg of pGag/Pol, 0.1µg of pREV, 0.1µg of pTAT, and 0.2µg of pVSVG) and 1.5µg of lentiviral vector. On days 3 and 4, the media was replenished with 3mL of fresh DMEM+glucose media. On days 4 and 5, the viral suspensions were harvested, pooled, pelleted at 1000g for 5min, and the supernatant was filtered through 0.45μm PES filters (Thermo, cat# 725-2545). The viral suspension was either used directly or kept frozen at −80°C until transduction. For transduction, U2OS and K562 cells were plated in 6-well plates (175000 and 200000 cells/well respectively) and infected with 0.5-2ml of viral suspension supplemented with polybrene at a final concentration of 8µg/mL. Infected cells were grown for several days before selection with antibiotics or FACS.

K562 dCas9-KRAB cells were previously published (Gilbert et al., 2013). U2OS dCas9-KRAB cells were generated by lentiviral transduction with pMH0006 (Addgene, cat# 135448; Chen et al., 2019) and selected for BFP expression by FACS. CRISPRi knockdown and control cell lines were generated by subsequent lentiviral transduction of dCas9 lines with plasmids containing individual sgRNAs (pOCIAD1sgRNA1 or pOCIAD1sgRNA2) or a non-targeting sgRNA and selected for higher levels of BFP expression by FACS. OCIAD1 knockdown cell lines rescued with wildtype or F102A OCIAD1 were generated by lentiviral transduction with plasmids pUltra-OCIAD1 and pUltra-OCIAD1(F102A), respectively, and selected for GFP expression by FACS. The U2OS OCIAD2 shRNA knockdown cells were generated by lentiviral transduction with plasmids containing individual shRNAs and selected with 15µg/ml blasticidin for 7 days. The U2OS OCIAD1/2 double knockdown cell line was generated by infecting stable U2OS CRISPRi cells stably expressing sgRNA#2 (above) with the lentivirus vector pLKO1-OCIAD2_shRNA1 and selecting infected cells with 15µg/ml blasticidin for 7 days. A control cell line was generated by infecting U2OS cells stably expressing a non-targeting sgRNA (above) with the lentivirus vector pLKO.1-blast-Scramble (Addgene, cat# 26701) expressing a non-targeting shRNA sequence and selected with 15µg/ml blasticidin for 7 days. Cell lines expressing truncated OCIAD1 constructs were generated by lentiviral infection of CRISPRi cells stably expressing sgRNA#2 (above) with the indicated pFUGW-OCIAD1 and pUltra-OCIAD1-TEV-StrepII lentiviral vectors. U2OS cells stably expressing matrix- or IMS-targeted GFP_1-10_ were generated by lentiviral transduction with the plasmids pMTS-GFP1-10 and pIMS-GFP1-10, respectively.

### Genome-scale CRISPRi screening

Genome-scale CRISPRi screens were conducted as previously described (Gilbert et al., 2014; Horlbeck et al., 2016; Jost et al., 2017). Briefly, K562 cells expressing dCas9-KRAB were transduced with the pooled hCRISPRi-v2 sgRNA library (Horlbeck et al., 2016) and selected for 2 days with 0.75µg/ml puromycin. Cells were then allowed to recover for 2 days in puromycin-free media before freezing library-containing cell aliquots (150×10^6^ cells per aliquot) under liquid nitrogen. After subsequent expansion and freezing while maintaining equivalent cell numbers, biological replicates were performed from two independent cell aliquots. Upon thawing, cells were recovered in RPMI+glucose for 4 days followed by 6 days conditioning in RPMI+galactose. At this point, t_0_ samples with a minimum 750x library coverage (150×10^6^ cells) were harvested while 250×10^6^ cells each were seeded in separate 3L spinner flasks (500ml of media at 0.5×10^6^ cells/ml) for treatment. Cells were treated with four pulses of antimycin (3.5-3.75nM) or vehicle (ethanol), consisting of 24h drug treatment, washout, and 48h recovery. For the duration of the screen, cells were maintained in RPMI+galactose at 0.5×10^6^ cells/mL by daily media dilution (minimum daily coverage approximately 1000 cells per sgRNA). At the end of the screen, endpoint samples from treated and vehicle-treated population (150×10^6^ cells each) were harvested and frozen. Genomic DNA was isolated from frozen cell pellets at the indicated time points and the sgRNA-encoding region was enriched, amplified, and processed for sequencing on an Illumina HiSeq 4000 platform as described previously (Horlbeck et al., 2016).

Sequencing reads were aligned to hCRISPRi-v2 library and counted using the Python-based ScreenProcessing pipeline (https://github.com/mhorlbeck/ScreenProcessing) (Horlbeck et al., 2016). Negative control genes were generated and phenotypes and Mann-Whitney *p*-values were calculated as described previously (Gilbert et al., 2014; Horlbeck et al., 2016; Jost et al., 2017). Briefly, antimycin A sensitivity phenotypes (ρ) were determined by calculating the log_2_ fold change in counts of an sgRNA in the treated and untreated samples, subtracting the equivalent median value for all non-targeting sgRNAs, and dividing by the number of population doubling differences between the treated and untreated populations (Gilbert et al., 2014; Jost et al., 2017; Kampmann et al., 2013). Phenotypes from sgRNAs targeting the same gene were collapsed into a single phenotype for each gene using the average of the three sgRNAs with the strongest phenotypes by absolute value and assigned a *p*-value using the Mann-Whitney test of all sgRNAs targeting the same gene compared to the non-targeting controls. For genes with multiple independent transcription start sites (TSSs) targeted by the sgRNA library, phenotypes and *p*-values were calculated independently for each TSS and then collapsed to a single score by selecting the TSS with the lowest Mann-Whitney *p*-value, as described previously (Gilbert et al., 2014; Horlbeck et al., 2016; Jost et al., 2017). Read counts and phenotypes for individual sgRNAs are available in source data 1. Gene-level phenotypes are available in source data 2.

### Validation of individual sgRNA phenotypes

The antimycin screen phenotype was validated by a growth competition assay using K562 cells expressing individually cloned sgRNAs. In short, K562 dCas9-KRAB cells were mixed with an equal number of K562 CRISPRi cells expressing a non-targeting sgRNA or sgRNA against OCIAD1. The sgRNA expression construct expressed a BFP reporter to identify infected cells. Of note, the dCas9-KRAB construct also expressed BFP fused to dCas9, but the BFP fluorescent intensity was dim and sgRNA-infected cells were clearly distinguishable from dCas9 cells by flow cytometry. For simplicity, K562 CRISPRi cells are refer to as BFP+ in the text. Cells were grown for 24h in RPMI+galactose containing either antimycin (5nM) or vehicle (ethanol), washed, and allowed to recover for 72h. The proportion of BFP-positive cells in each cell mixture was determined at the indicated time points using an Amnis Imagestream X (Luminex, Austin, TX) flow cytometer.

### Mitochondria isolation

For mitochondria isolation, all procedures were performed on ice or at 4°C. U2OS cells were grown to confluency in 150mm petri dishes and washed three times with 15mL of cold homogenization buffer (10mM HEPES, 1mM EDTA, 210mM mannitol, 70mM sucrose, pH 7.4 at 4°C). Cells were harvested by scraping in cold homogenization buffer (0.75mL per plate) supplemented with 1x protease inhibitor cocktail (MilliporeSigma, Burlington, MA) and lysed with 6-8 strokes of a glass Dounce homogenizer fitted with a tight pestle. At this point, a small fraction of homogenate was immediately snap frozen on liquid nitrogen and stored at −80°C for whole cell proteomics analysis as described below. For K562 suspension cells, cells were harvested by centrifugation (1000g, 5min), washed with cold homogenization buffer, re-pelleted (1000g, 5min), and incubated on ice for 20 min in swelling buffer (10mM HEPES, 1mM EDTA, pH 7.4 at 4°C) supplemented with 1x protease inhibitor cocktail (MilliporeSigma, Burlington, MA). Cells were then lysed with 25 strokes of a glass Dounce homogenizer fitted with a tight pestle and immediately diluted with 2x homogenization buffer (10mM HEPES, 1mM EDTA, 420mM mannitol, 140mM sucrose, supplemented with 1x protease inhibitor cocktail, pH 7.4 at 4°C) to a final concentration of 10mM HEPES, 1mM EDTA, 210mM mannitol, 70mM sucrose. The homogenate was centrifuged at ∼1300g for 5min to remove nuclei, unbroken cells, and large cellular debris and the supernatant was centrifuged at ∼14000g for 10min at 4oC. The crude mitochondrial pellet was resuspended in homogenization buffer supplemented with 1x protease inhibitor cocktail prior to measuring protein concentration using a bicinchoninic acid (BCA) assay (Pierce, Waltham, MA). Mitochondrial samples were either used immediately or snap frozen in 50 or 200µg aliquots on liquid nitrogen and stored at −80°C.

### Native PAGE analysis

Blue-Native (BN-) and Clear-Native (CN-) PAGE analyses were performed as previously described (Wittig et al., 2007, 2006). All procedures were performed on ice or at 4°C. Mitochondrial aliquots (200µg) were thawed on ice, diluted with 1mL of solubilization buffer (50mM imidazole, 50mM NaCl, 2mM 6-Aminohexanoic acid, 1mM EDTA, pH7.0 at 4°C), and pelleted at 21300g for 10min. The supernatant was removed and the mitochondrial pellet was resuspended in 20µL of solubilization buffer supplemented with digitonin (Calbiochem, cat# 300410) or Lauryl Maltose Neopentyl Glycol (LMNG) (Anatrace, Maumee, OH) to a final detergent-to-protein ratio of 4 g/g and 1 g/g, respectively. Samples were solubilized on ice for ∼15 min and centrifuged at 21300g for 20min. The supernatant was collected and protein concentration was measured using a BCA assay kit (Pierce, Waltham, MA).

For BN-PAGE, solubilized mitochondrial membranes were supplemented with 50% glycerol to a final concentration of 5% and Coomassie blue G-250 dye to a final detergent/dye ratio of 8 g/g. Equivalent amount of proteins were loaded on 3-12% polyacrylamide gels. The electrophoresis was started with cathode buffer B (50mM tricine, 7.5mM imidazole, 0.02% Coomassie blue G-250, pH ∼7.0) and exchanged with cathode buffer B/10 (50mM tricine, 7.5mM imidazole, 0.002% Coomassie blue G-250, pH ∼7.0) once the migration front had reached ∼1/3 of the resolving gel. For CN-PAGE, the solubilized mitochondrial samples were supplemented with 50% glycerol, 0.1% Ponceau S to a final concentration of ∼5% glycerol and ∼0.01% Ponceau S. Equivalent amount of proteins were loaded on 3-12% polyacrylamide gels. The cathode buffer contained 50mM tricine, 7.5mM imidazole, 0.01% dodecylmaltoside (DDM), and 0.05% sodium deoxycholate (DOC) (pH ∼7.0). The composition of the anode buffer (25mM imidazole, pH 7.0) was the same for BN-PAGE and CN-PAGE and remained constant for the duration of the electrophoresis. Gels were run in a cold room (4°C) at 100V until the samples had entered the resolving gel and at 275V thereafter. After electrophoresis, the gels were incubated in denaturing buffer (300mM Tris, 100mM acetic acid, 1% SDS, pH 8.6) at room temperature with agitation for 20 min and stored at room temperature between two glass plates for 1h to evenly distribute the SDS. Proteins were then electroblotted in at 4°C onto low fluorescent PVDF membranes at 90 mA and a voltage limited to 20V for 12-14h using a wet tank transfer apparatus filled with cold transfer buffer (150mM Tris, 50mM acetic acid, pH 8.6). BN-PAGE membranes were partially destained in 25% methanol, 10% acetic acid to visualize the ladder and completely destained with 100% methanol for Western blotting analysis. CN-PAGE membranes were stained with 5% acetic acid, 0.1% Ponceau S (w/v) to visualize the ladder and destained completely with extensive water washes before Western blotting analysis.

For 2D-native/SDS-PAGE analysis, individual gel lanes were excised from BN-PAGE gels immediately after electrophoresis and incubated in 8-10mL of denaturing buffer (62.5mM Tris pH 6.8, 2% SDS, 10% glycerol, 10mM TCEP) in a 15mL Falcon tube for 20 min at room temperature under gentle agitation. The gel strips were then equilibrated in 1x SDS-PAGE running buffer at room temperature for 15 min, loaded horizontally on a 10% polyacrylamide gel, and processed for Western blotting analysis as described below.

For the mobility shift assay, 400µg of K562 mitochondria was solubilized with LMNG at a 1g/g ratio as described above. The sample was halved and incubated with either mouse anti-PHB2 antibodies (Proteintech, cat# 66424-1-Ig, 70ng, ∼1.8µl) or vehicle (PBS) on ice of 90min. Samples were then analyzed by BN-PAGE as described above.

### Protease protection and carbonate extraction analysis

Protease protection analysis was performed on mitochondria freshly isolated from U2OS cells as previously described (Hoppins et al., 2011) with the following modifications. Mitochondria (50µg of total mitochondrial protein) were resuspended in 500µl of one of the following solutions: homogenization buffer (210mM mannitol, 70mM sucrose, 10mM HEPES, 1mM EDTA, pH 7.4), mitoplast/swelling buffer (10mM HEPES, pH 7.4), or solubilizing buffer (homogenization buffer with 1% Triton X-100). After 15 min incubation on ice, the mitoplast/swelling sample was gently pipetted up and down 15 times to disrupt the outer mitochondrial membrane. Proteinase K was then added to the indicated samples to a final concentration of 100µg/mL, and samples were incubated on ice for 20 min. The digestion was stopped by adding PMSF to a final concentration of 2mM and incubating the samples on ice for 5 min. TCA was then added to a final concentration of 12.5% and proteins were precipitated on ice for 1h. Proteins were then pelleted by centrifugation at 21130g for 15 min at 4°C, washed with acetone, dried, and resuspended in 100µl of 1x Laemmli buffer. Samples (20µl) were loaded on a 10% SDS-PAGE and analyzed by Western blotting with the indicated antibodies as described below.

The carbonate extraction assay was performed as described (Hoppins et al., 2011) with the following modifications. Mitochondria isolated from U2OS cells (50µg of total mitochondrial protein) were thawed on ice, pelleted at 15000g for 10 min at 4°C, and resuspended in 200µl of one of the following solutions: 10mM HEPES (pH 7.4), 100mM sodium carbonate (pH 10.5), 100mM sodium carbonate (pH 11), or 100mM sodium carbonate (pH 11.5). Samples were incubated on ice for 30 min and centrifuged at 100000g for 1h in a TLA100 rotor. The supernatant was harvested and proteins were precipitated with TCA as described above. The pellet fraction and TCA-precipitated proteins were resuspended in 50µl of 1x Laemmli buffer and 10µl was loaded on a 10% SDS-PAGE and analyzed by Western blotting with the indicated antibodies as described below.

### Western blotting analysis

For quantitative Western blot analysis, protein concentration was determined using a BCA assay kit (Pierce, Waltham, MA) and equivalent amount of proteins were diluted with 6x Laemmli sample buffer to a final concentration of 62.5mM Tris pH 6.8, 2% SDS, 10% glycerol, 0.1M DTT, 0.01% bromophenol blue. Samples were heated for 2-5min at 95°C and loaded on 10% Tris-glycine polyacrylamide gels. After electrophoresis, proteins were electroblotted on low fluorescent PVDF or nitrocellulose membranes, and immunoblotted with the following primary antibodies: rabbit anti-OCIAD1 (Invitrogen, cat# PA5-20834, 1:2000-1:5000), mouse anti-OCIAD1 (Proteintech, cat# 66698-1-Ig, 1:5000), rabbit anti-OCIAD2 (Invitrogen, cat# PA5-59375, 1:500-1:5000), mouse anti-ATP5A1 (Proteintech, cat# 66037-1-Ig, 1:2000-1:5000), rabbit anti-NDUFB8 (Proteintech, cat# 14794-1-AP, 1:2000), mouse anti-SDHA (SantaCruz Biotechnology, cat# sc-166947, 1:2000-1:5000), rabbit anti-UQCRC2 (Proteintech, cat# 14742-1-AP, 1:2000-1:5000), mouse anti-UQCRC1 (Invitrogen, cat# 459140, 1:2000), rabbit anti-CYC1 (Proteintech, cat# 10242-1-AP, 1:1000), mouse anti-COXIV (Proteintech, cat# 66110-1-1g, 1:2000), mouse anti-PHB2 (Proteintech, cat# 66424-1-Ig, 1:5000), rabbit anti-TIM50 (Proteintech, cat# 22229-1-AP, 1:1000), rabbit anti-TOM70 (Proteintech, cat# 14528-1-AP, 1:1000), mouse anti-GFP (Proteintech, cat# 66002-1-Ig, 1:2000), mouse anti-β-actin (Proteintech, cat# 66009-1-1g, 1:10000). Secondary antibodies conjugated to DyLight 680 and DyLight 800 (Thermo Fisher Scientific, 1:5000) were used and visualized with an Odyssey Infrared Imaging System (LI-COR, Lincoln, NE). Densitometry analysis was done using the quantification software ImageStudio Lite (LI-COR, Lincoln, NE).

### Heme detection

Chemiluminescence was used to detect c-type heme on PVDF or nitrocellulose membranes as previously described (Dorward, 1993; Feissner et al., 2003). In short, membranes were rinsed with distilled water immediately after electrophoresis, incubated with SuperSignal West Femto chemiluminescent substrate (Pierce, Waltham, MA), and imaged on an ImageQuant LAS 4000 (GE, Boston, MA). Densitometry analysis was done using the quantification software ImageStudio Lite (LI-COR; Lincoln, NE).

### GFP complementation assay

U2OS cells stably expressing GFP_1-10_ in the matrix (MTS) or IMS were plated in 6-wells plate (∼300000 cells/well) on day 0 and transfected on day 1 with 6µl of Lipofectamine 2000 (Invitrogen, Carlsbad, CA), according to the manufacturer’s instructions. The cells were transfected with 250ng of the following plasmids: CoQ9-GFP_11_, GFP_11_-MICU1, OCIAD1-GFP_11_, and GFP_11_-OCIAD1, and 750ng of transfection carrier DNA (Promega, pGEM2 plasmid). Cells were expanded in 10cm plate on day 2 and analyzed by fluorescent flow cytometry on day 3 with an Amnis Imagestream X (Luminex, Austin, TX).

### Immunopurification

Cells were crosslinked with dithiobis(succinimidyl propionate) (DSP, Life Technologies, cat# 22585) made from a freshly prepared 0.25M stock solution in DMSO. In short, 150ml of confluent (∼1×10^^^6 cells/ml) K562 cells of the indicated OCIAD1 background were harvested by centrifugation (1000g, 5min), washed with warm (37°C) PBS, and crosslinked at room temperature for 30 min with 0.5mM DSP in PBS at ∼1×10^^^6 cells/ml. DSP was then quenched by adding Tris-HCl (pH 7.5) to a final concentration of 100mM. Cells were harvested by centrifugation (1000g, 5 min), washed with cold PBS, harvested again, and solubilized in 2ml of cold RIPA buffer supplemented with 1x protease inhibitor cocktail (MilliporeSigma, Burlington, MA) on ice for 30 min. Samples were centrifuged at 26000g for 30 min at 4°C in a TLA100.4 rotor. The supernatant was collected, protein concentration was measured using a BCA assay kit (Pierce, Waltham, MA), and aliquots were stored at −80°C.

Immunopurification was performed on three independently DSP-crosslinked samples. Each sample was thawed on ice and adjusted to 7.8mg of total protein in 2ml of RIPA buffer containing 1x protease inhibitor cocktail (MilliporeSigma, Burlington, MA). OCIAD1 was immunocaptured overnight at 4°C with 3µg of rabbit anti-OCIAD1 antibody (Thermo Fisher, cat# PA5-20834). Antibodies were captured with 100µl of μMACS protein A beads (Miltenyi Biotec; San Diego, CA). Beads were isolated with μ columns and a μMACS separator (Miltenyi Biotec; San Diego, CA), washed 5 times with 1ml of RIPA buffer and 3 times with 1ml of 50mM ammonium bicarbonate pH 8.0. Bait proteins were eluted with 25µl of elution buffer (2M Urea, 0.67M thiourea in 50mM Ammonium bicarbonate pH 8.0) containing LysC/Trypsin (Promega, Madison, WI, cat# V5071) to a final concentration of 5µg/ml followed by two elution with 50µl of elution buffer without LysC/Trypsin. Samples were reduced with 10mM TCEP (Pierce, Waltham, MA) for 30 min at 37°C, alkylated with 15mM 2-chloroacetamide (MilliporeSigma, Burlington, MA), digested overnight at 37°C, and desalted using ZipTip with 0.6 µL C18 resin (MilliporeSigma, Burlington, MA, cat# ZTC18S096) prior to LC-MS/MS analysis as described below.

### Protein digestion on suspension traps

Protein digestion of U2OS lysates was done on suspension traps (S-Trap) as described (Ludwig et al., 2018) with the following modifications. Whole cell and crude mitochondrial lysates (50µg total protein) were boiled in 5% SDS, 50 mM ammonium bicarbonate (pH 7.55) for 5 min. Proteins were then reduced with 10mM TCEP for 15 min at 37°C and alkylated in the dark for 30 min with 15 mM 2-chloroacetamide. The proteins were then acidified with phosphoric acid (final concentration of 1.2%) and diluted with 6 volumes of S-Trap buffer (90% methanol, 100 mM ammonium bicarbonate, pH 7.1). The colloidal suspension was loaded onto DNA miniprep spin columns used as “suspension traps” (EZ-10 DNA spin columns, Biobasic, Amherst, NY) and washed with S-Trap buffer prior to overnight proteolysis at 37°C with LysC/trypsin (Promega, Madison, WI) in 50 mM ammonium bicarbonate (pH 8.0) at a protease/protein ratio of 1:40 (w/w). Peptides were successively eluted with 40µl of 50 mM ammonium bicarbonate (pH 8.0), 40µl of ultrapure Milli-Q water, 0.1% TFA, and 40µl of 80% acetonitrile, 0.1% TFA in ultrapure Milli-Q water. Peptides were dried using a SpeedVac concentrator and resuspended in 30µl of 2% acetonitrile, 0.1% TFA. Peptide concentration was measured using a fluorometric peptide assay kit (Pierce, Waltham, MA) and samples were analyzed by LC-MS/MS as described below.

### In-gel protein digestion

To minimize contamination, procedures were performed in a biosafety cabinet whenever possible. Mitochondria from U2OS cells of the indicated OCIAD1 background were solubilized with digitonin at a 4g/g detergent/protein ratio and 100µg of solubilized mitochondrial protein was resolved by BN-PAGE as described above. After electrophoresis, the gel was fixed with 40% methanol, 10% acetic acid at room temperature for 20 min and destained with 8% acetic acid for 20 min. Gel slices (2mm x 7mm) were excised along the entire lane using disposable gel cutter grids (The Gel Company, San Francisco, CA, cat# MEE2-7-25). Ten gel slices ranging from ∼600-900kDa were collected in 100µl of 50mM ammonium bicarbonate (pH 8.0) in a 96-well plate and destained/dehydrated with successive 5 min washes with 100µl of the following solutions (3 washes each): 50mM ammonium bicarbonate (pH 8.0), 25% acetonitrile in 50mM ammonium bicarbonate (pH 8.0), 50% acetonitrile in 50mM ammonium bicarbonate (pH 8.0), 75% acetonitrile in 50mM ammonium bicarbonate (pH 8.0), 100% acetonitrile. Proteins were then reduced with 50µl of 10mM TCEP for 30 min at 37°C, gel slices were dehydrated again with three washes with 100% acetonitrile, and alkylated with 15mM 2-chloroacetamide in the dark for 20 min. Gel slices were dehydrated again and washed for 5 min with 100µl of the following solutions (2 washes each): 50mM ammonium bicarbonate (pH 8.0), 25% acetonitrile in 50mM ammonium bicarbonate (pH 8.0), 50% acetonitrile in 50mM ammonium bicarbonate (pH 8.0), 75% acetonitrile in 50mM ammonium bicarbonate (pH 8.0) and four washes with 100% acetonitrile. Gel slices were air-dried before overnight ProteaseMax-aided digestion as previously described (Saveliev et al., 2013). In short, dried gel pieces were rehydrated in 50µl of 12 ng/µl LysC/Trypsin (Promega, Madison, WI), 0.01% ProteaseMAX surfactant (Promega, Madison, WI, cat# V2071) in 50mM ammonium bicarbonate (pH 8.0) for 20 min on ice and overlaid with 50µl of 0.01% ProteaseMAX surfactant in 50mM ammonium bicarbonate (pH 8.0). Proteins were digested overnight at 37°C. The peptide-containing solution was collected in 1.5ml eppendorf tubes and 100µl of 75% acetonitrile, 1% TFA in 25mM ammonium bicarbonate (pH 8.0) was added to each gel slice to elute remaining peptides. Both eluates were pooled and dried using a SpeedVac concentrator before LC-MS/MS analysis as described below.

### Mass spectrometry analysis

LC-MS/MS analysis was performed at the University of California, Davis, Genome Center Proteomics Core. Immunoprecipitation and whole cell samples were run on a Thermo Scientific Fusion Lumos mass spectrometer in Data Independent Acquisition (DIA) mode. Peptides were separated on an Easy-spray 100µm x 25cm C18 column using a Dionex Ultimate 3000 nUPLC with 0.1% formic acid (solvent A) and 100% acetonitrile, 0.1% formic acid (solvent B) and the following gradient conditions: 2% to 50% solvent B over 60 minutes, followed by a 50%-99% solvent B in 6 minutes, held for 3 minutes and finally 99% to 2% solvent B in 2 minutes. The total run time was 90 minutes. Six gas phase fractionated (GPF) chromatogram library injections were acquired using 4Da staggered isolation windows (GPF 1: 400-500 m/z, GPF 2: 500-600 m/z, GPF 3: 600-700 m/z, GPF 4: 700-800 m/z, GPF 5: 800-900 m/z, and GPF 6: 900-1000 m/z). Mass spectra were acquired using a collision energy of 35, resolution of 30K, maximum inject time of 54 ms and a AGC target of 50K. The analytical samples were run in DIA mode with 8 Da staggered isolation windows covering 400-1000 m/z.

BN-PAGE gel samples were run on a Bruker TimsTof Pro mass spectrometer. Peptides were directly loaded on a Ionoptiks (Parkville, Victoria, Australia) 75µm x 25cm 1.6µm C18 Aurora column with Captive Spray emitter. Peptides were separated using a Bruker Nano-elute nUPLC at 400nl/min with 0.1% formic acid (solvent A) and 100% acetonitrile, 0.1% formic acid (solvent B) and the following gradient conditions: 2% solvent B to 35% solvent B over 30min. Runs were acquired in diaPASEF mode (Meier et al., 2020) with an acquisition scheme consisting of four 25 m/z precursor windows per 100ms TIMS scan. Sixteen TIMS scans, creating 64 total windows, layered the doubly and triply charged peptides on the m/z and ion mobility plane. Precursor windows began at 400 m/z and continued to 1200 m/z. The collision energy was ramped linearly as a function of ion mobility from 63 eV at 1/K0=1.5 Vs cm−2 to 17 eV at 1/K0=0.55 Vs cm−2.

Raw files acquired in DIA mode on the Fusion/Lumos instrument were analyzed with DIA-NN 1.7.12 (Demichev et al., 2020) using the following settings (Protease: Trypsin/P, Missed cleavages: 1, Variable modifications: 0, Peptide length range: 7-30, Precursor m/z range: 300-1800, Fragment ion m/z range: 200-1800, Precursor FDR: 1). The N-term M excision, C carbamidomethylation, M oxidation, and RT profiling options were enabled and all other parameters were set to default. To generate a sample-specific spectral library, we initially used DIA-NN to create a large proteome-scale *in silico* deep learning-based library from the Uniprot human reference proteome (UP000005640, one protein per gene) with a list of common contaminants. This large spectral library was refined with deep sample specific chromatogram libraries. In short, equal amount of peptides from all U2OS cell lines (control, OCIAD1 knockdown, OCIAD2 knockdown, OCIAD1/2 double knockdown, and OCIAD1 knockdown rescued with wildtype OCIAD1) were pooled to create a master sample containing all peptides theoretically identifiable within our samples. To maximize the depth of our library, whole cell lysate and mitochondrial pooled samples were processed separately. Deep chromatogram libraries were created from these pooled samples using six gas-phase fractionated DIA injections with a total of 52 overlapping 4 m/z-wide windows ranging from 400 to 1000m/z as previously described (Searle et al., 2018). The resulting chromatogram libraries were used together with the large predicted deep learning-based spectral library to generate a new highly optimized spectral library. This new spectral library was subsequently used to process our analytical samples.

Raw files acquired in diaPASEF mode on the timsTOF were analyzed similarly with DIA-NN (version 1.7.13 beta 1) using the following settings (Protease: Trypsin/P, Missed cleavages: 1, Variable modifications: 0, Peptide length range: 7-30, Precursor m/z range: 300-1800, Fragment ion m/z range: 200-1800, Precursor FDR: 1, MS1 mass accuracy: 10ppm, MS2 mass accuracy: 10ppm). The N-term M excision, C carbamidomethylation, and M oxidation options were enabled and all other parameters were set to default. In short, we generated a deep learning-based predicted library from the Uniprot human reference proteome (UP000005640, one protein per gene) supplemented with N-terminal truncated CYC1 isoforms and a list of common contaminants. This large library was used to process all raw files from the gel slices analytical runs and generate a second and more optimized spectral library that includes ion mobility data. This new highly optimized spectral library was finally used to re-analyze all raw files.

DIA-NN output files were imported and analyzed in R using MaxLFQ values quantified from proteotypic peptides only (Cox et al., 2014). For whole cell proteomics and immunoprecipitation analysis, only proteins identified in at least all the replicates of a given sample were selected. Missing values were imputed using the “MinDet” deterministic minimal value approach from the MSnbase package prior statistical analysis as described below.

### Confocal fluorescence microscopy

U2OS cells were grown on 12mm round glass coverslips (#1.5) and stained for 30 min with 100nM of Mitotracker DeepRed (Invitrogen, Carlsbad, CA, cat# M22426), washed with PBS, and fixed in 4% PFA in PBS for 20 min at room temperature. Cells were washed again with PBS, permeabilized for 10 min with 0.1% Triton X-100 in PBS, blocked with 5% bovine serum albumine (BSA) in PBS for 1h at room temperature, and immunolabeled with rabbit anti-OCIAD1 (Invitrogen, cat# PA5-20834, 1:10000) or rabbit anti-OCIAD2 (Invitrogen, cat# PA5-59375, 1:5000) antibodies for 1h at room temperature in 1% BSA in PBS. Cells were washed again in PBS and incubated with donkey anti-rabbit IgG conjugated with AlexaFluor 488 (Invitrogen, Carlsbad, CA, cat# A21206, 1:1000) in 1% BSA in PBS for 1h at room temperature. Finally, cells were washed again in PBS and mounted on glass slides with ProLong Glass antifade mounting medium (Invitrogen, Carlsbad, CA, cat# P36980). Images were collected using the spinning-disk module of a Marianas SDC Real Time 3D Confocoal-TIRF microscope (Intelligent Imaging Innovations; Denver, CO) fitted with a 100×, 1.46 NA objective and a Hamamatsu (Japan) Orca Flash 4.0 sCMOS camera. Images were captured with SlideBook (Intelligent Imaging Innovations) and linear adjustments were made using ImageJ.

### Multiple sequence alignment

Multiple sequence alignment analysis was performed with the R package “msa” (version 1.22.0) using the Clustal Omega method with default parameters.

### Statistical analysis

Gene ontology (GO) enrichment analysis was performed using the topGO R package (version 2.42.0) using the ‘elim’ method and Fisher’s exact test (Alexa et al., 2006; Grossmann et al., 2007). Western blot densitometry results were analyzed using one-way analysis of variance (ANOVA) followed by pairwise t-test with Benjamini & Hochberg (FDR) correction. For LC-MS/MS immunoprecipitation and whole cell proteomics data, relative changes between conditions were analyzed using limma’s function lmFit (Ritchie et al., 2015), followed by eBayes with false-discovery rate correction (Phipson et al., 2016). For whole cell proteomics data, hierarchical clustering was performed using Euclidean distances of significant hit proteins. Error bars represent standard error and *p < 0.05, **p < 0.01, and ***p < 0.001. All data were analyzed in R (version 4.0.3).

## Acknowledgments

We would like to thank Vadim Demichev for helping with DIA-NN analysis. We also want to thank James A. Letts and María Maldonado for their valuable feedback on this manuscript. FACS sorting at UC Davis is supported by the NIH (S100D018223). MJ is supported by funding from the NIH (grant K99 GM130964). JRF is supported by funding from the NIH (R00HL133372 and R35GM137894) and the Welch Foundation (I-1951-20180324). Min Y Cho coordinated resource sharing between labs and assisted with screen sample processing.

---

**Figure 1-figure supplement 1.**
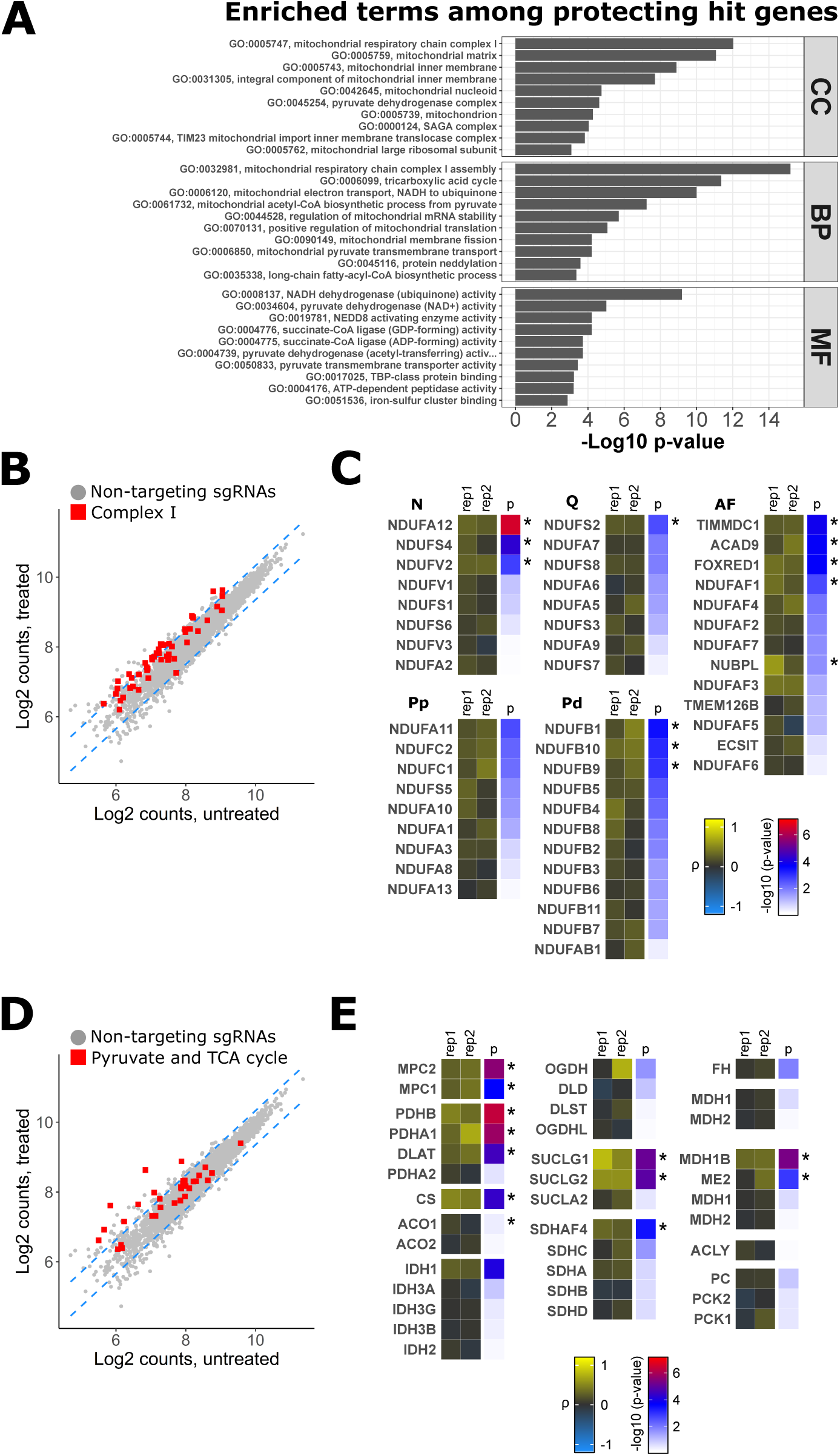
Silencing genes related to Complex I, pyruvate, and TCA metabolism protects cells against chemical inhibition of Complex III. A) Top 10 categories from gene ontology (GO) enrichment analysis for protecting hit genes (ρ > 0). Mitochondrial terms related to Complex I, pyruvate metabolism, and the TCA cycle are specifically enriched in all three biological domains (CC: Cellular Component, BP: Biological Process, MF: Molecular Function). Terms are ordered by the maximum of -log10 p-value (elim algorithm with Fisher’s exact test). B,D) Read count distribution of non-targeting control sgRNAs (grey circles) and sgRNAs related to Complex I structural subunits and assembly factors (B) or sgRNAs related to pyruvate metabolism and TCA cycle (D) in untreated and antimycin treated cells. Red squares represent the average read count of the top 3 targeting sgRNAs. Dashed blue lines represent the 95% prediction interval. C,E) Tile plots displaying the phenotype scores (ρ) (first and middle columns) and associated p-values (right column) of both biological replicates for complex I related genes (C) and genes related to pyruvate and TCA metabolism (E). Complex I genes were grouped by module (N = N-module; Q = Q-module; Pp = proximal peripheral arm; Pd = distal peripheral arm) and assembly factors (AF). Significant genes are indicated by asterisks.

**Figure 1-figure supplement 2.**
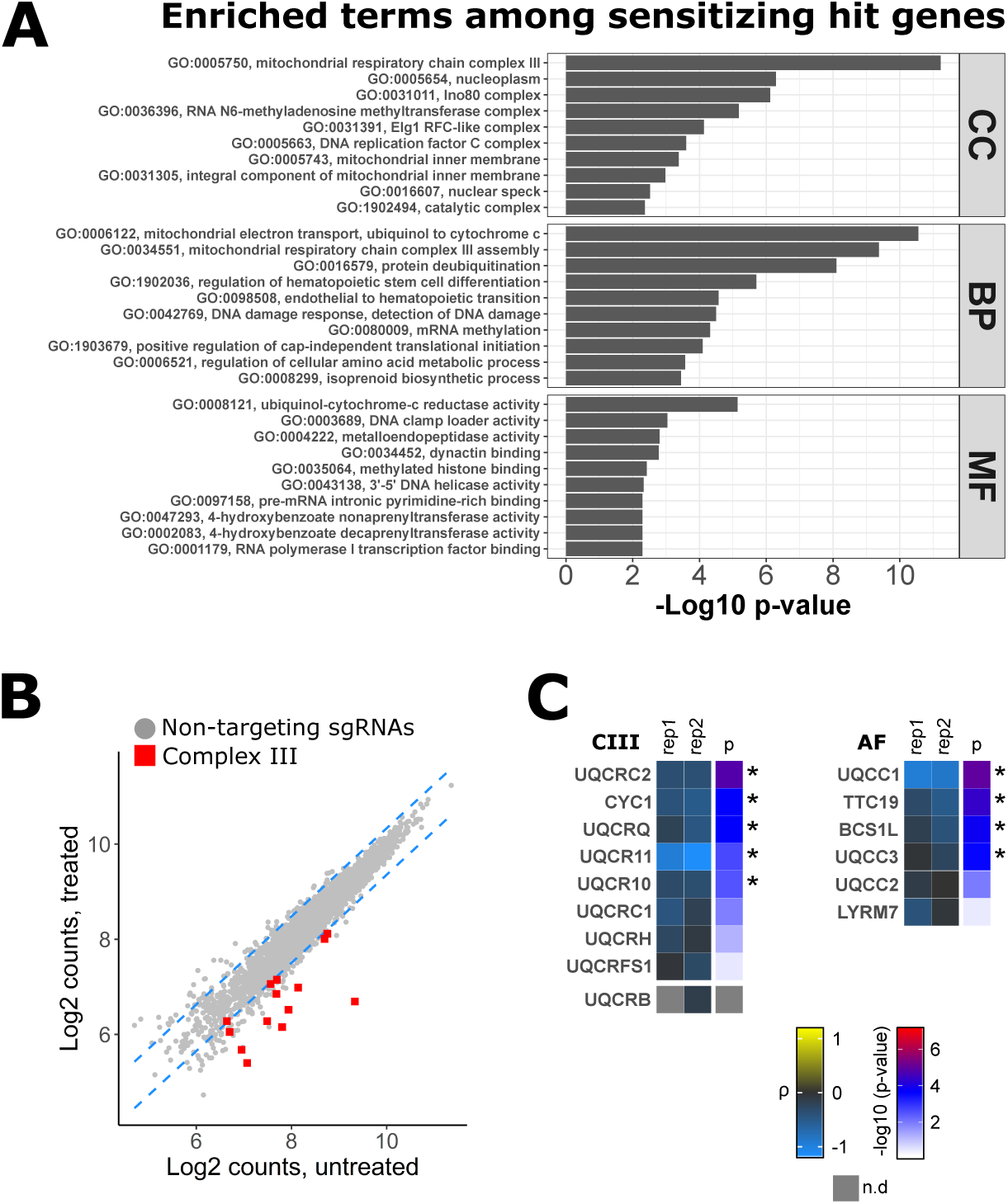
Silencing Complex III genes aggravate the cellular response to antimycin A. A) Sensitizing antimycin A genes. Top 10 categories from gene ontology (GO) enrichment analysis (ρ < 0). Mitochondrial terms related to Complex III are enriched in all three biological domains (CC: Cellular Component, BP: Biological Process, MF: Molecular Function). Terms are ordered by the maximum of -log10 p-value (elim algorithm with Fisher’s exact test). B) Read count distribution of non-targeting control sgRNAs (grey circles) and Complex III structural subunit and assembly factor sgRNAs (req squares, average of top 3 sgRNAs) in untreated and antimycin treated cells. Dashed blue lines represent the 95% prediction interval. C) Tile plots displaying the phenotype scores (ρ) of each biological replicate (first and middle columns) and associated p-values (right column) for CIII_2_ structural genes (CIII) and assembly factors (AF). Significant genes are indicated by asterisks. (n.d = not determined).

**Figure 3-figure supplement 1.**
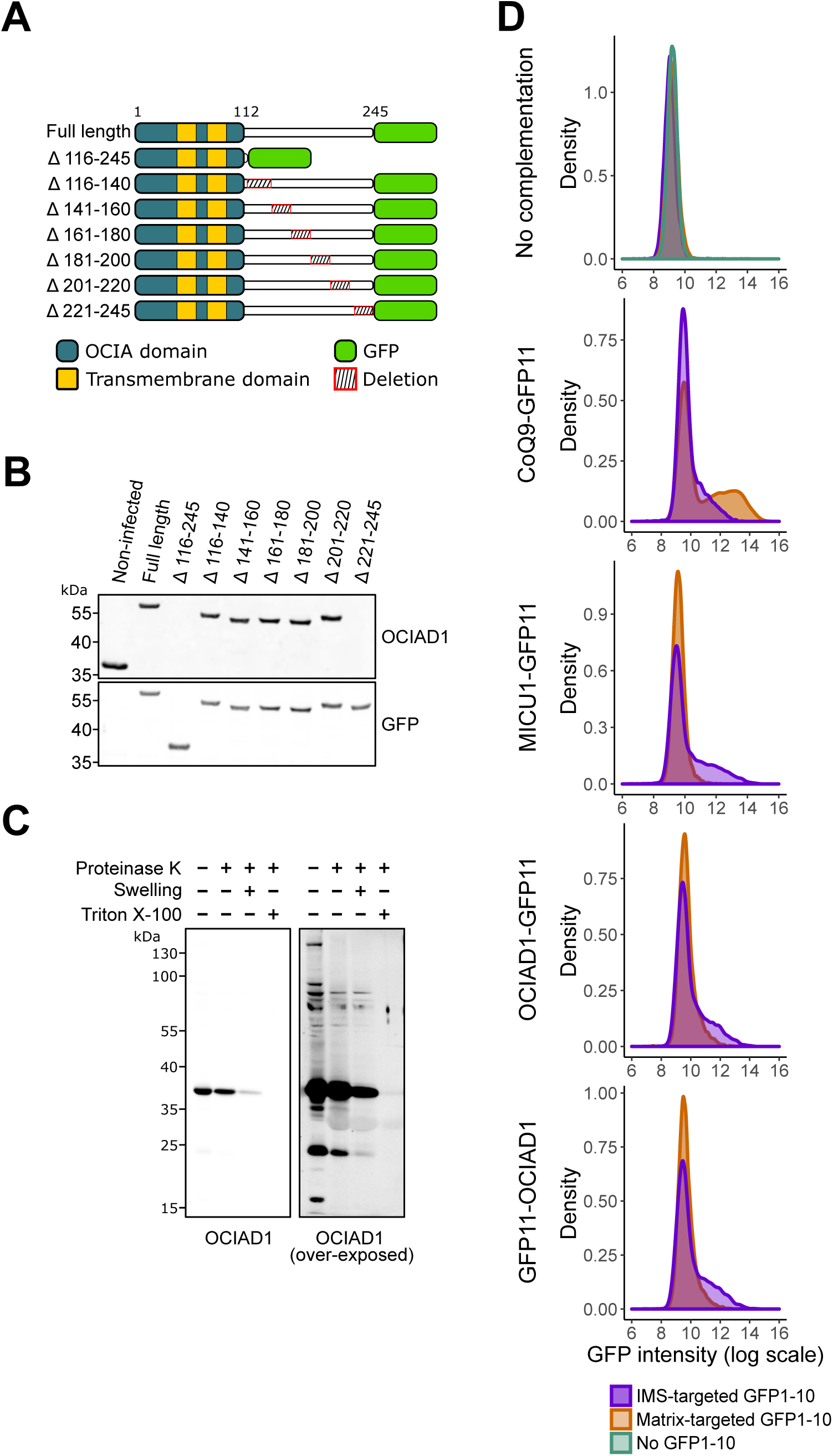
OCIAD1 termini are localized in the mitochondrial intermembrane space. A) Schematic illustrating the OCIAD1-GFP deletion constructs used for mapping the epitope of the anti-OCIAD1 polyclonal antibody. B) Cell lysates from U2OS cells expressing either full-length or truncated OCIAD1-GFP were analyzed by Western blotting and immunoprobed using anti-OCIAD1 (Invitrogen, cat# PA5-20834) and anti-GFP antibodies. The anti-OCIAD1 polyclonal antibody recognizes an epitope located within the last 25 amino acids of OCIAD1 C-terminus. C) Uncropped immunoblot for the OCIAD1 protease protection assay shown in Figure 3C alongside an over-exposed image of the same membrane. D) U2OS cells stably expressing IMS- or matrix-targeted GFP_1-10_ were transiently transfected with various GFP_11_ constructs before assessing GFP complementation by flow cytometry analysis. No GFP_1-10_ and GFP_1-10_ alone (uppermost panel). CoQ9 tagged with C-terminal GFP_11_ expressed in matrix- or IMS-targeted GFP_1-10_ cells (second panel). MICUI tagged with C-terminal GFP_11_ expressed in matrix- or IMS-targeted GFP_1-10_ cells (third panel). N-terminal GFP_11_-tagged OCIAD1 construct (fourth panel) and C-terminal GFP_11_- tagged OCIAD1 construct (fifth panel) expressed in matrix- or IMS-targeted GFP_1-10_ cells.

**Figure 4-figure supplement 1.**
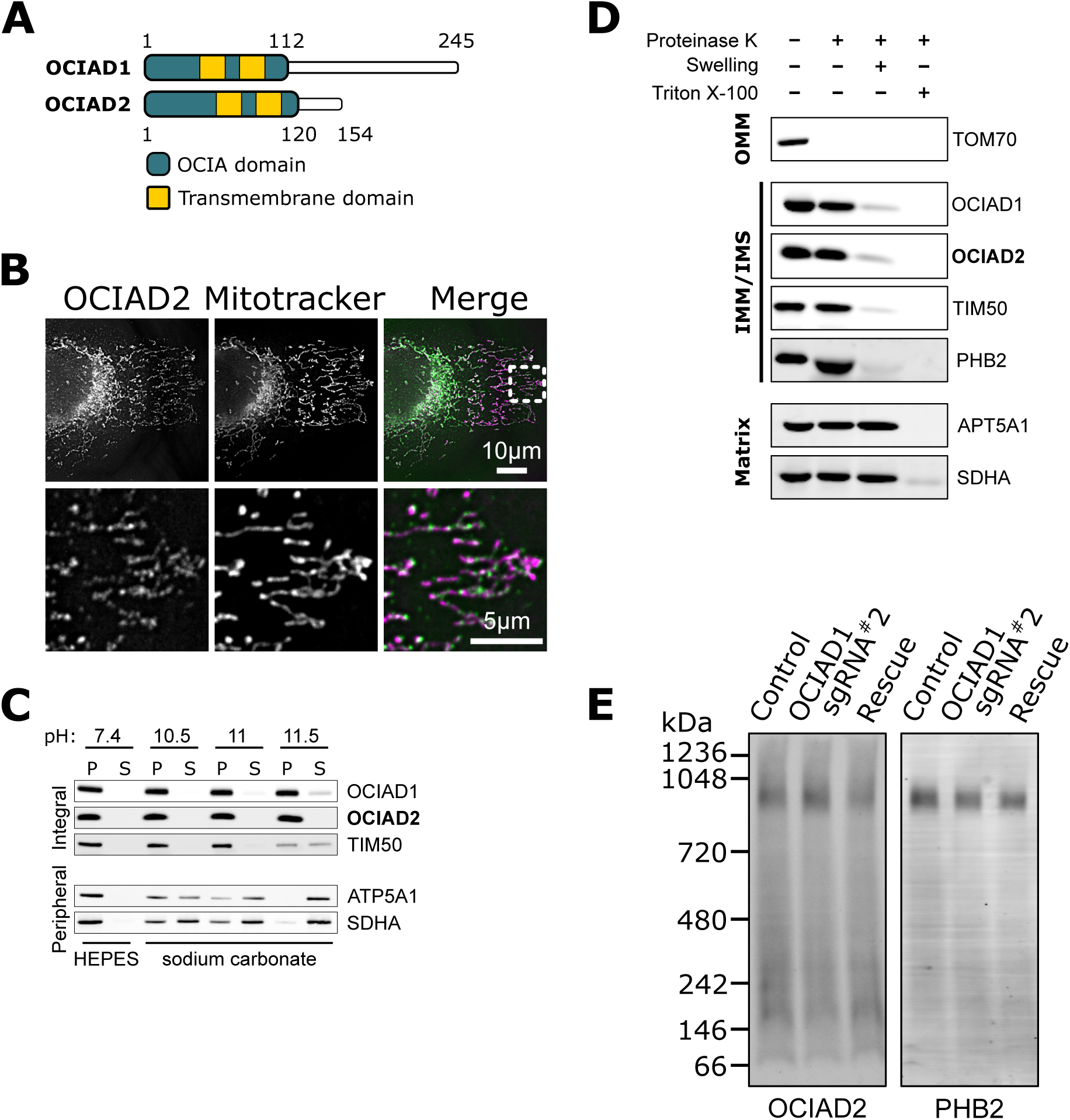
The OCIAD1 paralog, OCIAD2, localizes to the mitochondria inner membrane. A) Schematic of predicted OCIAD1 and OCIAD2 topologies. B) Representative images of fixed U2OS cells stained with Mitotracker (magenta) and immunolabeled for OCIAD2 (green). The bottom panel is a magnification of the inset shown in the upper panel. C) OCIAD2 is an integral membrane protein. Sodium carbonate extraction fractions (pH 10.5-11.5) immunoblotted with anti-OCIAD1, anti-OCIAD2, anti-TIM50, anti-ATP5A1, and anti-SDHA antibodies. P and S indicate pellet and soluble fractions, respectively. This panel, without the OCIAD2 blot, was shown in Figure 3C. E) OCIAD2 localizes to the inner membrane. Protease protection assay fractions immunoblotted with anti-OCIAD1, anti-OCIAD2, anti-prohibitin 2 (PHB2), anti-TIM50, anti-ATP5A1, and anti-SDHA antibodies. (OMM: outer mitochondrial membrane, IMM: inner mitochondrial membrane, IMS: intermembrane space). This panel, without the OCIAD2 blot, was shown in Figure 3D. F) BN-PAGE of LMNG detergent-solubilized mitochondrial membranes isolated from U2OS cells and immunoblotted with anti-OCIAD2 and anti-prohibitin antibodies.

**Figure 4-figure supplement 2.**
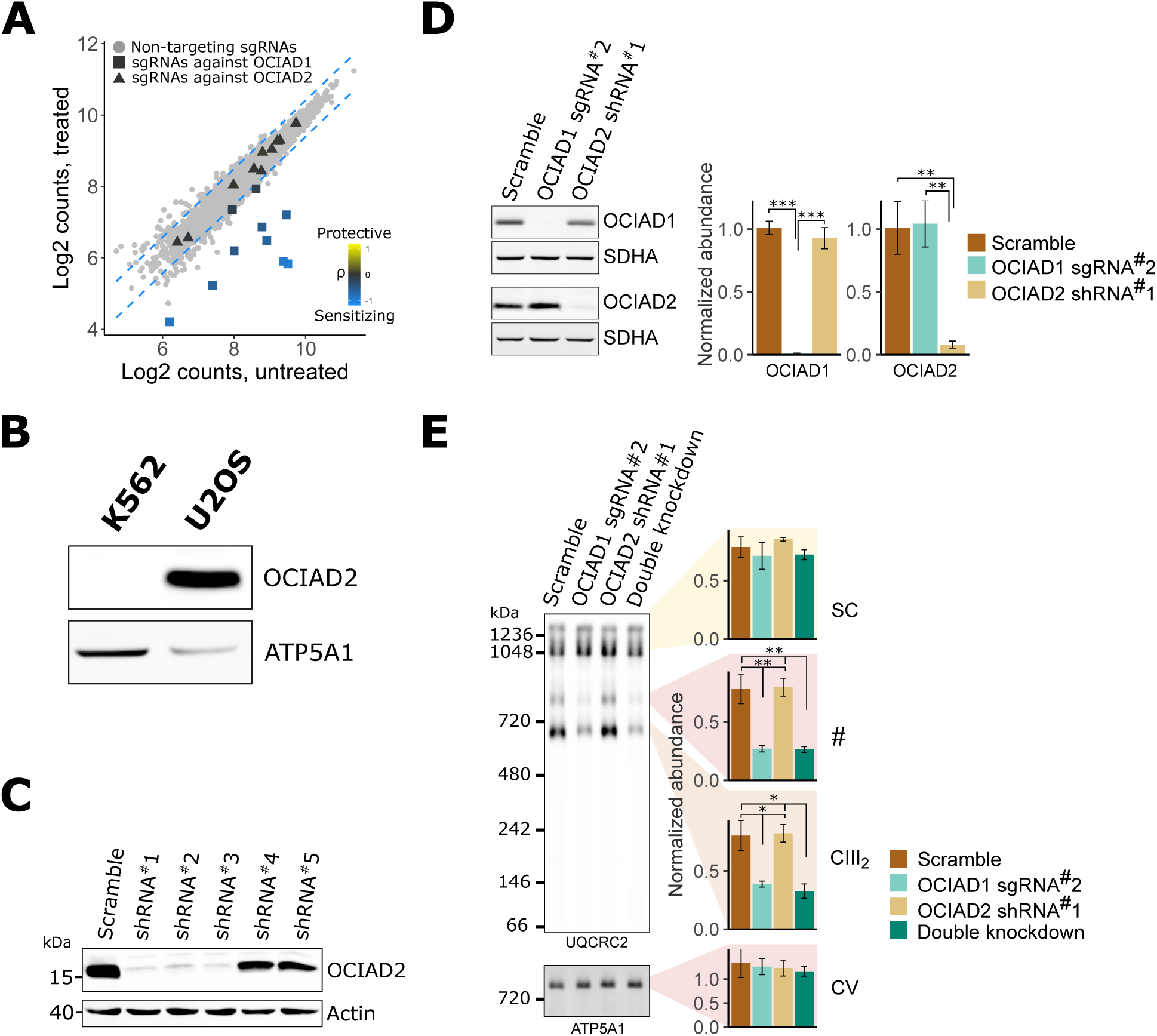
OCIAD1 and OCIAD2 paralogs are functionally divergent. A) Read count distribution of all 10 sgRNAs targeting OCIAD1 (squares) and OCIAD2 (triangles) in untreated and antimycin treated K562 cells. Grey circles represent non-targeting sgRNAs. Dashed blue lines represent the 95% prediction interval. B) Western blot of U2OS and K562 cell extracts immunoblotted with anti-OCIAD2 and anti-ATP5A1 antibodies. C) Western blot of cell extracts from U2OS cells expressing scramble or OCIAD2 shRNAs. The membrane was immunoblotted with anti-OCIAD2 and anti-β-actin antibodies. D) Western blot of extracts from U2OS cells expressing OCIAD2 shRNA and OCIAD1gRNA. The membrane was immunoblotted with anti-OCIAD2 and anti-SDHA antibodies. SDHA was used as a loading control. E) BN-PAGE analysis of digitonin-solubilized mitochondrial extracts from U2OS cells expressing OCIAD1 sgRNA#2 and OCIAD2 shRNA#1. ATP5A1 served as a loading control. Values represent normalized intensity ± SEM (n = 3). Asterisks (*p < 0.05, **p < 0.01, or ***p < 0.001) correspond to the adjusted (FDR) p-values from the post-ANOVA pairwise t-test.

**Figure 4-figure supplement 3.**
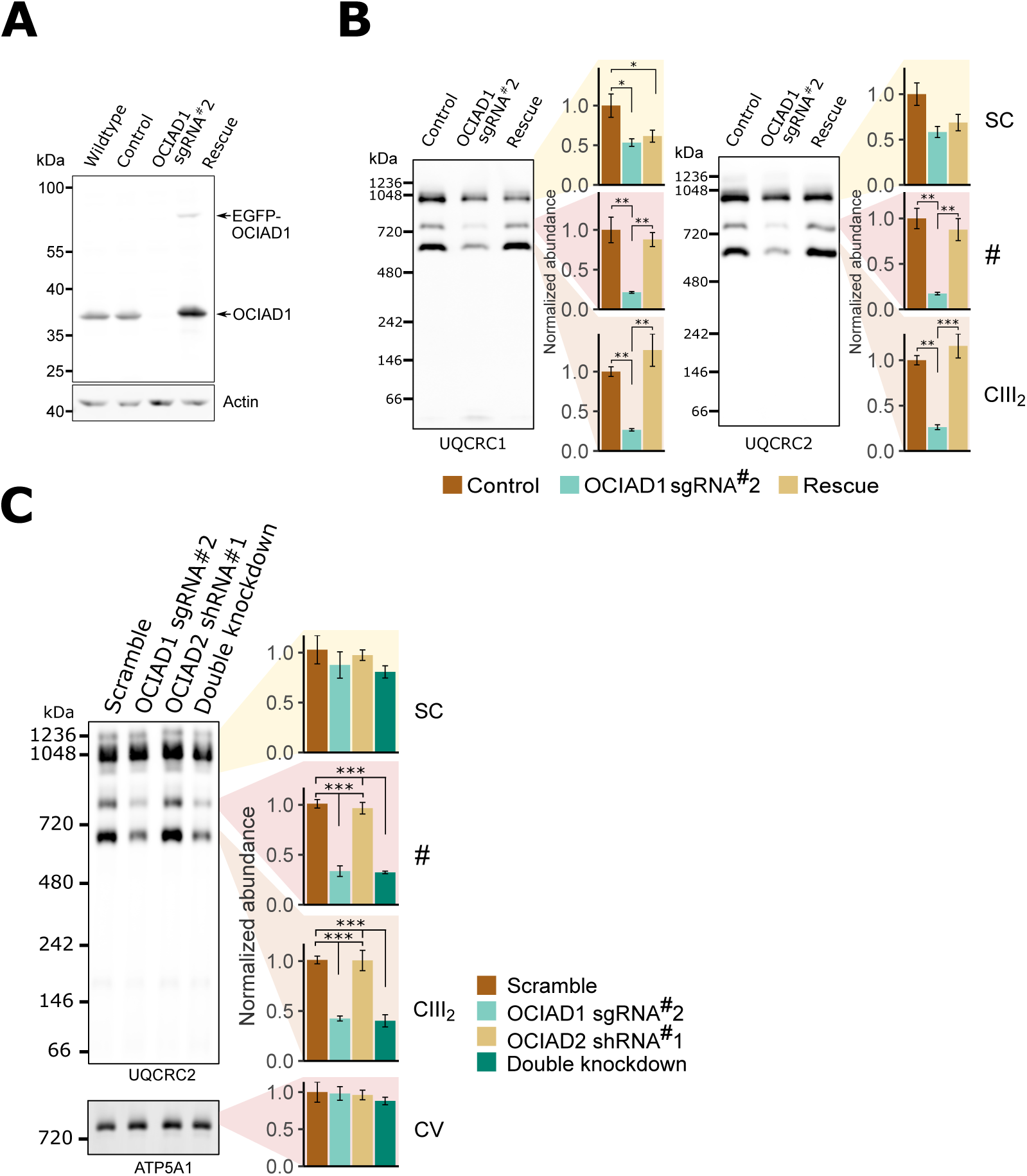
The role of OCIAD1 in CIII2 assembly is independent of cell-type and glucose availability. A) Western blot of wildtype U2OS cells or U2OS cells expressing a non-targeting sgRNA, sgRNA#2 against OCIAD1, or sgRNA#2 and wildtype OCIAD1 rescued by lentivirus expression. The upper band (EGFP-OCIAD1) represents intact fusion gene product. The non-targeting sgRNA used in this study does not affect OCIAD1 expression (compare control and wildtype lanes). B) BN-PAGE results using two CIII_2_ core subunits (UQCRC1, left and UQCRC2, right) showing that OCIAD1 is also required for CIII_2_ assembly in U2OS cells grown in glucose-containing media. C) BN-PAGE indicating that silencing OCIAD1, but not OCIAD2, disrupts CIII_2_ assembly in U2OS cells grown in glucose-containing media. ATP5A1 served as a loading control. Values represent normalized intensity ± SEM (n = 3). Asterisks (*p < 0.05, **p < 0.01, or ***p < 0.001) correspond to the adjusted (FDR) p-values from the post-ANOVA pairwise t-test.

**Figure 4-figure supplement 4.**
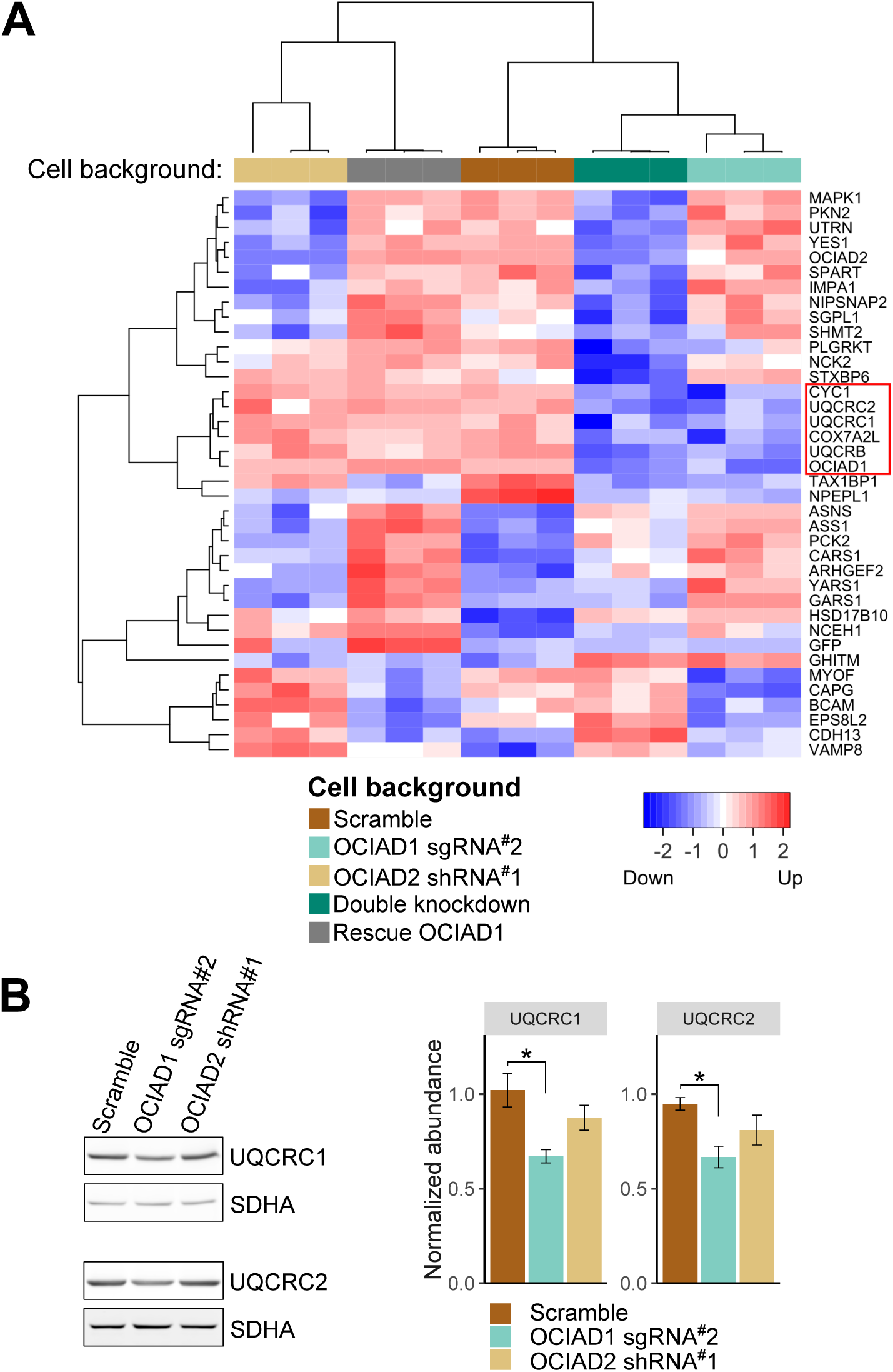
OCIAD1 regulates steady-state levels of Complex III subunits. A) Hierarchical clustering of unbiased proteomic analysis performed on whole-cell lysate from U2OS control cells, OCIAD1 knockdown cells, OCIAD2 knockdown cells, OCIAD1 and OCIAD2 double knockdown cells, and OCIAD1 knockdown cells rescued with wildtype OCIAD1. The analysis identified a small cluster (red square) enriched for Complex III proteins selectively down-regulated in the OCIAD1 and OCIAD1/OCIAD2 knockdown cells. B) Western blot analysis showing that two Complex III subunits (UQCRC1 and UQCRC2) are downregulated in mitochondria isolated from OCIAD1 knockdown U2OS cells but not from OCIAD2 knockdown U2OS cells. SDHA served as a loading control.

**Figure 4-figure supplement 5.**
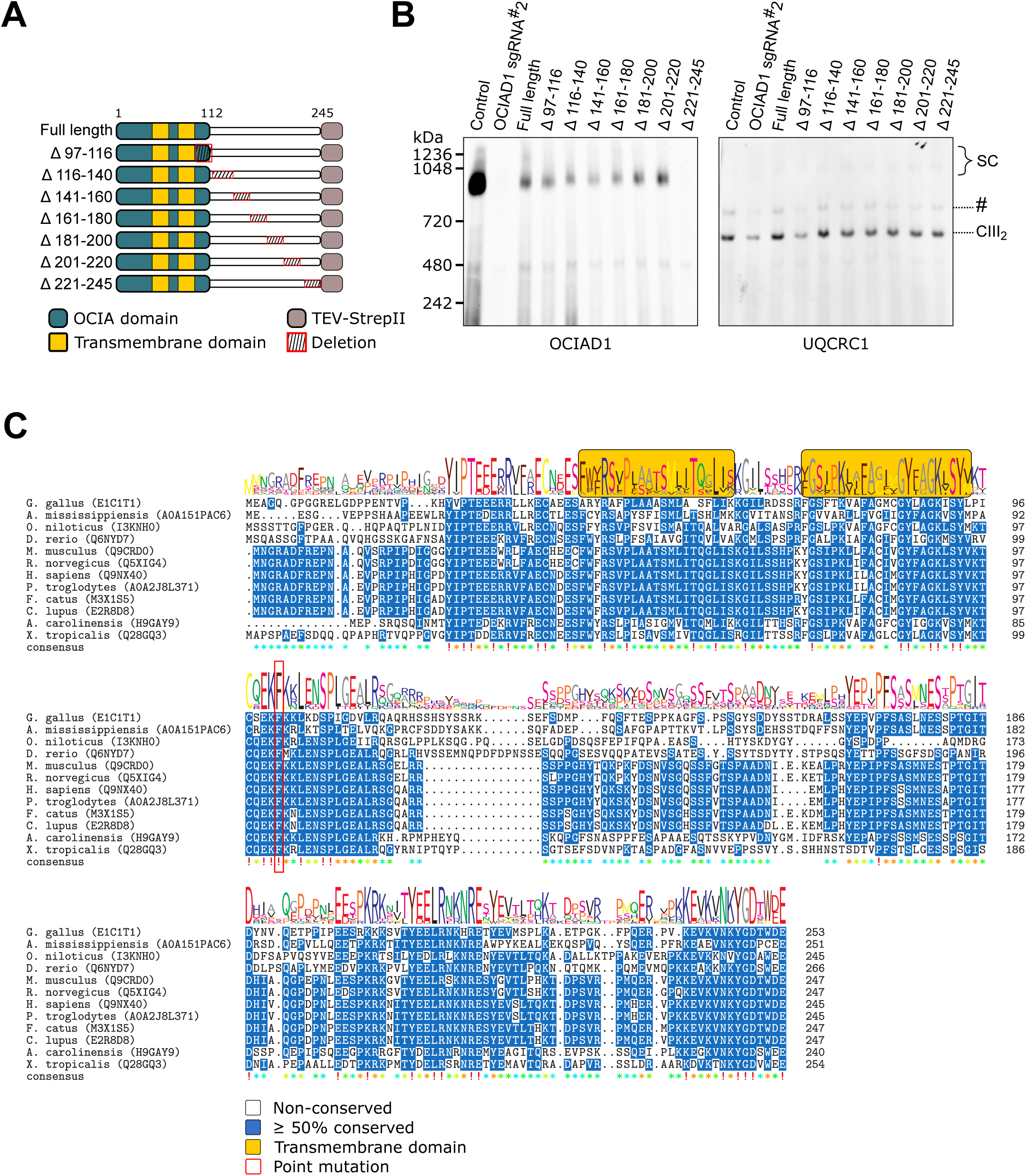
The distal region of the OCIA domain is essential for the function of OCIAD1 in CIII_2_ assembly. A) Schematic showing the topology of the OCIAD1-TEV-StrepII isoforms with C-terminal tiled deletions. B) Mitochondria were isolated from non-infected control and OCIAD1 knockdown cells, or OCIAD1 knockdown cells infected with a lentivirus expressing full-length or truncated OCIAD1 isoforms. BN-PAGE followed by Western blot analysis for OCIAD1 and UQCRC1 identified a small portion of the OCIA domain (a.a. 97-116) as putatively essential for CIII_2_ assembly. C) Multiple sequence alignment of OCIAD1 protein sequences using Clustal Omega. Blue shading indicates over 50% of identical amino acids in all sequences. The red box indicates the location of the mutated residue.

**Figure 5-figure supplement 1.**
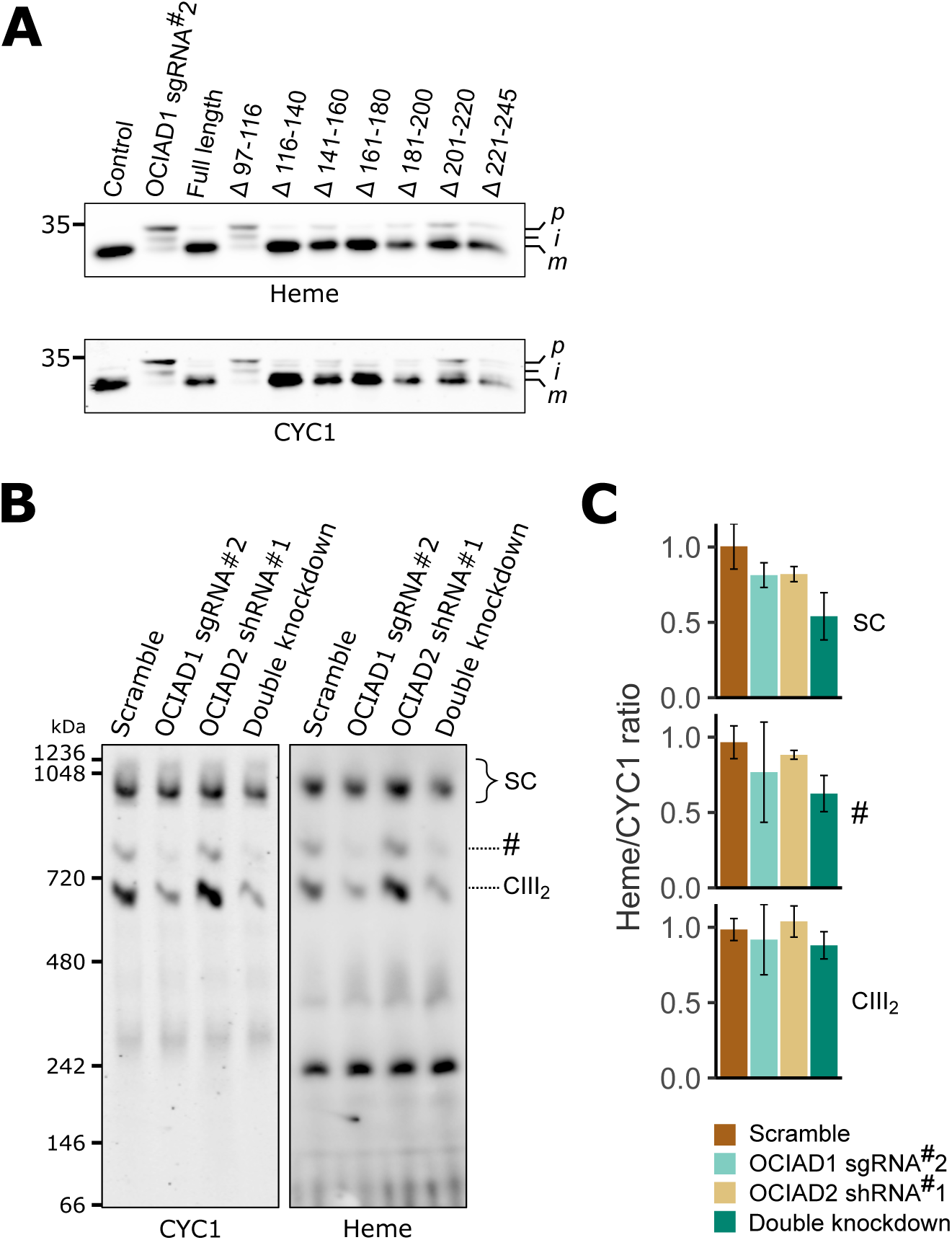
Mature CIII_2_ contains hemylated cytochrome *c_1_* in OCIAD1 knockdown cells. A) Western blot analysis of mitochondria isolated from OCIAD1 knockdown cells rescued with truncated OCIAD1 isoforms shown in Supplementary Figure 8 A and B. Heme was detected by chemiluminescence before immunoblotting the membrane with an antibody against CYC1. Deleting the distal portion of the OCIA domain (Δ97-116) disrupted CYC1 maturation. B) Blue-native PAGE analysis of digitonin-solubilized mitochondrial membranes isolated from U2OS control and OCIAD1 knockdown cells. Heme was detected by chemiluminescence before immunoblotting the membrane with an antibody against Complex III core subunit UQCRC2. C) Clear-native PAGE analysis of digitonin-solubilized mitochondrial membranes isolated from U2OS control and OCIAD1 knockdown cells. Heme was detected by chemiluminescence before immunoblotting the membrane with an antibody against CYC1. D) Quantification of blots shown in B and C showing the proportion of hemylated CYC1 in CIII_2_ assemblies.

